# Primate thalamic nuclei select abstract rules and shape prefrontal dynamics

**DOI:** 10.1101/2024.03.13.584871

**Authors:** Jessica M. Phillips, Mohsen Afrasiabi, Niranjan A. Kambi, Michelle J. Redinbaugh, Summer Steely, Emily R. Johnson, Xi Cheng, Maath Fayyad, Sounak Mohanta, Asia Carís, Charles B. Mikell, Sima Mofakham, Yuri B. Saalmann

## Abstract

Flexible behavior depends on abstract rules to generalize beyond specific instances, and outcome monitoring to adjust actions. Cortical circuits are posited to read out rules from high-dimensional representations of task-relevant variables in prefrontal cortex (PFC). We instead hypothesized that converging inputs from PFC, directly or via basal ganglia (BG), enable thalamus to select rules. We measured activity across PFC and connected thalamic nuclei of monkeys applying rules. Abstract rule information first appeared in ventroanterior thalamus (VA) – the main thalamic hub between BG and PFC. Mediodorsal thalamus (MD) also represented rule information before PFC, persisting to help maintain activation of relevant PFC cell ensembles. MD, a major recipient of midbrain dopamine input, was first to represent information about behavioral outcomes. A PFC-BG-thalamus model reproduced key findings, and thalamic-lesion modeling disrupted PFC rule representations. This suggests that thalamus selects high-level cognitive information from PFC and monitors behavioral outcomes of these selections.

## INTRODUCTION

Cognitive control is our ability to flexibly adapt behavior according to goals and context ^1,2^. It can be considered hierarchical in the sense that goals can be selected at different levels of abstraction ^3,4^. Prefrontal cortex (PFC) is vital for cognitive control. PFC neurons respond to different task variables (mixed selectivity ^5^) and adapt to current task demands (adaptive coding ^6^), to form a large, flexible workspace. These high-dimensional representations allow readout of task-relevant rules, to either guide behavior, or to be fed back to PFC neurons to influence processing of upcoming task events for hierarchical cognitive control ^5,7^. Although downstream cortical neurons have been posited to perform this readout, convergent PFC projections to the thalamus directly, or via the basal ganglia (BG), could enable thalamic readout, which would subsequently be disseminated via divergent thalamo-PFC projections. This would enable strengthening of initially weak PFC signals, updating of PFC representations and adaptive coding.

The PFC is extensively interconnected with several nuclei in the medial thalamus ^8^. There are two main types of pathways issued from the PFC which synapse onto thalamocortical neurons ^9^ (Fig. 1G). Projections from cortical layer 5 (Fig. 1G, warm colored corticothalamic projections) drive thalamocortical neurons which preferentially target PFC layer 1 (Fig. 1G, orange thalamocortical projections). These thalamocortical neurons have a modulatory influence on PFC, and project back to the PFC in both a reciprocal (i.e., forming “local” transthalamic loops) and unreciprocated (i.e., forming “long-range” transthalamic loops) manner ^8^. These thalamocortical neurons often stain positively for the calcium binding protein calbindin and have been referred to as “matrix”-type neurons ^10^ (although there is some heterogeneity among “matrix” neurons ^11^). Such thalamic matrix neurons are thought to play an important role in modulating cortical gain and establishing contextually appropriate functional connectivity ^8,12,13^. Output from cortical layer 6 (Fig. 1G, green corticothalamic projections), on the other hand, modulates thalamocortical neurons which preferentially target PFC middle layers (Fig 1G, green thalamocortical projections). These thalamocortical neurons have a driving influence on PFC, and project back to PFC in a reciprocal manner (i.e., forming “local” transthalamic loops) ^8^. These thalamocortical neurons stain positively for the calcium binding protein parvalbumin and are known as “core”-type neurons ^10^. Their projections to cortical middle layers may help sustain cortical activity when feedforward sensory input is removed. Of the different thalamic nuclei that connect with PFC, two stand out as possible key players in hierarchical cognitive control required for rule-guided behavior: the mediodorsal thalamic nucleus (MD) and ventroanterior thalamic nucleus (VA) ^8^.

**Figure 1.**
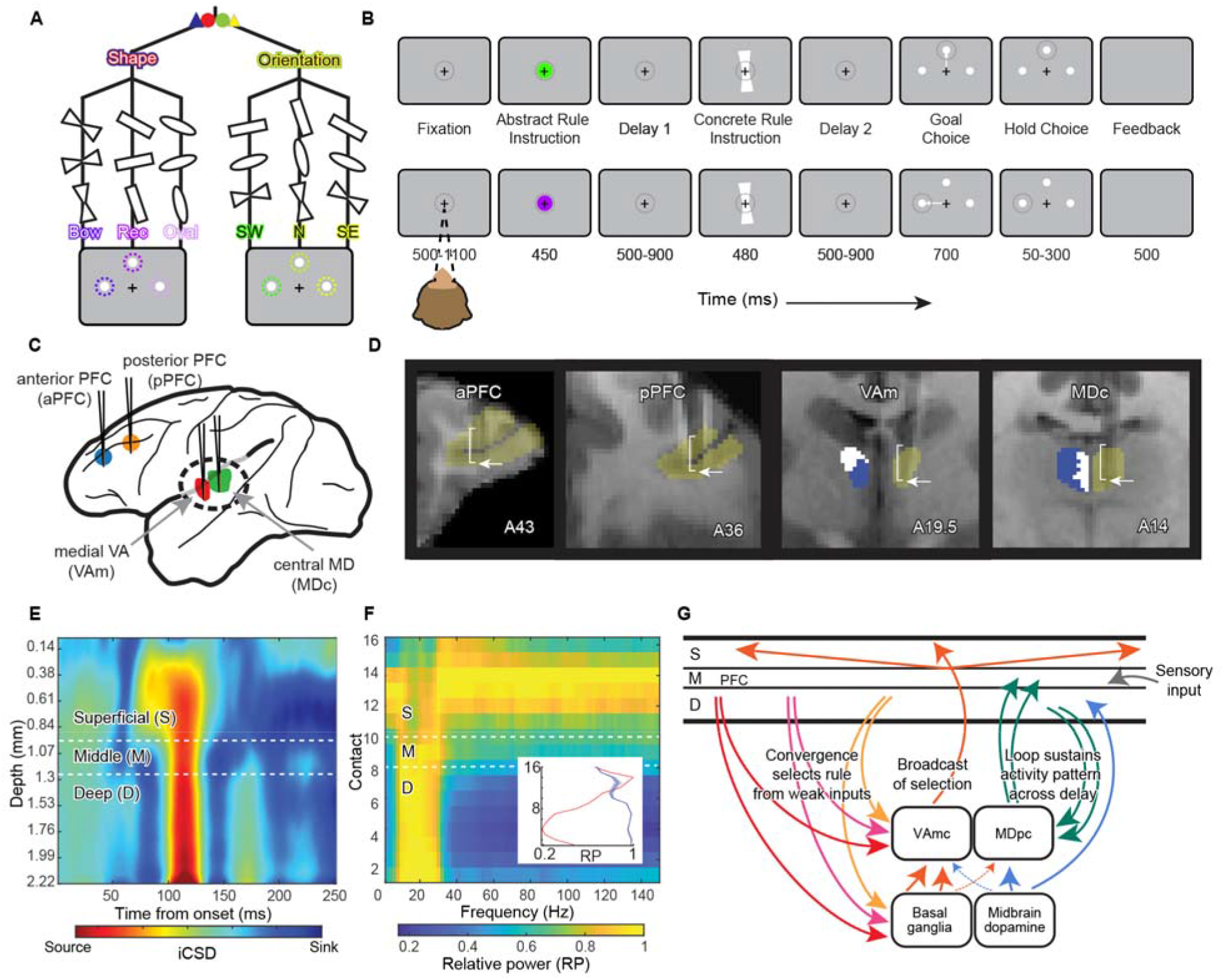
Experimental design and hypotheses. (**A**) HRT stimulus permutations. There were two abstract rules (shape, orientation) each specifying a set of concrete rules (i.e., cues mapping onto action/targets). (**B**) HRT trial structure. Two trials with identical concrete rule stimuli but different preceding abstract rule instructions require distinct stimulus-response mappings to obtain reward. Although we used blue and red cues to instruct shape, and green and yellow cues to instruct orientation, in (B) (and elsewhere) we represent shape instruction using purple and orientation instruction using yellowish green. In top row, abstract cue instructs that orientation is relevant, and because the concrete rule following delay 1 is oriented “north”, monkeys select vertical target (see A, fifth column). In bottom row, abstract cue instructs that shape is relevant, and because this same concrete cue is a bowtie shape, monkeys select leftward target (see A, first column). (**C**) Lateral view of monkey brain showing location of ROIs. (**D**) Coronal structural MRIs showing probes *in situ*. Blue voxels show SNr projection zones in VAm and PFC projection zones in MDc (only displayed for left hemisphere, to allow probe visualization in right hemisphere), obtained using probabilistic tractography. Bracket indicates span of LMA contacts (∼4.6 mm) and arrow indicates probe tip. (**E** and **F**) Calculation of CSD and relative LFP power map enabled designation of LMA contacts as situated within superficial, middle and deep layers. Inset in F shows relative power (RP) averaged in the alpha-beta (blue) and gamma (red) frequency band as a function of contact number, i.e., laminar depth ^23^. (**G**) Projections to and from VAmc and MDpc with PFC layers. Inputs from the mostly deep layer 5 of multiple PFC architectonic regions (warm colored corticostriatal and corticothalamic arrows) issue convergent driving inputs in VAmc and the striatum (input nucleus of BG). Converging PFC inputs funneled through BG influence gating of thalamocortical neurons in VAmc by the SNr. When disinhibited by SNr neurons, VAmc thalamocortical neurons (matrix) can issue widespread, modulatory signals to superficial layers. Inputs from mostly deep layer 6 of more restricted PFC zones issue modulatory projections to MD thalamocortical neurons. Impacted MD core neurons return driving inputs to middle layers of same PFC zone (local, reciprocal). Ratio of matrix to core neurons is approximately 50/50 for VAmc and 20/80 for MDpc. Bow – bowtie; Rec – rectangle; SW – southwest; N – north; SE – southeast; RP – relative power.

The MD is well known for its close anatomical relationship with the PFC ^10^. The parvocellular compartment (MDpc) has the strongest relationship with the dorsolateral PFC ^10^, important for rule-guided behavior ^1^. MDpc contains mostly “core” thalamocortical neurons innervated by cortical layer 6, which form “local” transthalamic loops and have a driving influence on PFC middle layers ^8,10^ (Fig. 1). In mice, such loops reinforce PFC representations, important for working memory processes ^14,15^. But mouse MD neurons show little/no rule information and sustain PFC representations via a nonspecific volley of activity ^15^. This may be related to the reduced cognitive capacity in mice and relatively low numbers of rule-selective neurons in mouse PFC (compared with primates) ^16^. In primates, MD neurons are known to work closely with lateral PFC to process precise visual and motor information important for behavior guided by spatial working memory ^17^. One electrophysiology study also suggested that MD neurons may pass go/nogo rule-related information to the PFC, although the neuron sample size was small (n=7) ^18^. Another species difference is that, unlike mice, primate MD receives major dopaminergic input ^19^, suggesting an additional outcome monitoring role may be possible ^20-22^. This is consistent with observations that thalamic lesion patients (where MD is affected) show reduced amplitude of the error-related negativity and behavior consistent with impaired performance monitoring ^22^. We therefore hypothesized that the contributions of primate MD to cognitive control are not limited to processing and maintenance of visuospatial information, rather that MD also encodes other key behaviorally relevant information such as the (1) currently relevant rule and (2) behavioral outcome. We also predicted that MD neurons would carry information about these task variables across extended windows of time, having a more general role sustaining activity in cortical circuits for information maintenance and processing.

Only found in primates ^10,24^, the VA magnocellular compartment (VAmc) is anatomically connected with most if not all PFC regions, similar to MD ^8^. PFC inputs converge quite massively in the VAmc ^25^, similar to the cortico-striatal projection system ^26^. Unlike MDpc, VAmc contains more “matrix” neurons innervated by cortical layer 5, which participate in long-range transthalamic paths and have a modulatory influence in PFC superficial layers ^8,10^ (Fig. 1G). VAmc thalamocortical neurons are the main target of BG output issued from the substantia nigra pars reticulata (SNr) and appear to be tightly controlled by its tonic inhibitory influences ^27^. The SNr is the main BG output node that issues information to the PFC ^27,28^. This may include rule-related information, which has been reported to be represented in a subpopulation of striatal neurons ^29,30^, and a preliminary sample of VAmc neurons ^18^. Thus, firing patterns of VAmc neurons likely reflect changes in BG outflow relevant for PFC function. Unlike MD neurons which have a more focal influence, the influence of VAmc neurons appears to be more widespread, impacting an assortment of PFC architectonic regions simultaneously ^18^. We hypothesized that multiple levels of convergence from PFC to VAmc, either directly or via the BG (which itself is characterized by convergent input patterns ^31^), enable VAmc neurons to select task-relevant rules. Because VAmc is equipped with a suite of efferent projections that spread across multiple PFC regions, we propose that VAmc facilitates the initiation of task sets, by modulating cortical gain, recruiting PFC neurons to active ensembles representing rules and establishing optimal functional connectivity ^8^, for processing of incoming lower-level stimuli based on higher-level rules.

## RESULTS

### Diffusion MRI-guided electrophysiological recordings

Two monkeys learned the hierarchical rule task (HRT), requiring the sequential application of two rules (Fig. 1, A and B, and fig. S1). It is hierarchical in the sense that the initial abstract rule cue (e.g., green circle symbolizing orientation set) specifies a class of subordinate, more concrete rules (i.e., southwest (SW)-oriented cue mapping onto leftward target, north (N)-oriented cue mapping onto vertical target, southeast (SE)-oriented cue mapping onto rightward target), similar to tasks used with humans ^32,33^. As such, the concrete rule (unlike the abstract rule) can be mapped onto a motor response. Further, abstract rules are defined as those which can be generalized across similar but novel situations to guide appropriate behavior, and indeed the monkey behavior was consistent with abstract rule generalization (i.e., across new irrelevant stimulus features during training (fig. S1)). While monkeys performed the HRT, we used linear microelectrode arrays (LMAs) to simultaneously record electrophysiological activity from four cortical and subcortical regions of interest (ROIs; Fig. 1, C and D). For PFC recording sites, we targeted the principal sulcus (putative areas 46d/v) in two locations separated by approximately 7 mm in the anterior-posterior plane, thus yielding anterior PFC (aPFC) and posterior PFC (pPFC) sites in each monkey. We used diffusion MRI and probabilistic tractography to target locations in MD and VA with a high likelihood of connectivity with the PFC sites (Fig. 1D, blue voxels). For VA, we additionally targeted voxels which had a high probability of connection with the SNr, indicating the likely location of VAmc (Fig. 1D, A19.5 panel, blue voxels). Thus, our thalamic LMAs were situated in central MD (MDc, putative MDpc) and medial VA (VAm, putative VAmc). We use these more conservative terms and abbreviations (i.e., MDc and VAm) throughout the rest of this article when referring to our recording sites, to distinguish them from the architectonic subnuclei that we targeted, which require use of specific staining approaches with post-mortem tissue for their precise definition.

We measured spiking activity in a total of 1094 neurons in four ROIs (229 in aPFC, 340 in pPFC, 196 in VAm, 329 in MDc; all included in whole-ROI analyses). In cortex, we used current source density ^34^ (CSD; Fig. 1E) and unipolar local field potential (LFP) analyses ^23,35^ (Fig. 1F) to designate electrode contacts and neurons to cortical layers (n = 147 superficial, n = 104 middle, n = 272 deep; all included in layer-specific analyses; remaining PFC cells could not be assigned to a particular layer due to partial laminar span in that recording session). Of the aPFC and pPFC neurons, we designated 65 and 82 to superficial, 54 and 50 to middle and 105 and 167 to deep layers, respectively.

### Thalamus selects abstract rules

Many thalamic neurons showed robust, short-latency selectivity for the currently relevant abstract rule (Fig. 2, A to C, E to G and fig. S2, A to F). We used two different methods to quantify selectivity of neurons. First, we used a relatively conservative nonparametric test to compare the number of spikes across conditions, which suggested a subset in each population was rule-selective (aPFC, 2.6%, fig. S2G; pPFC, 10.3%, Fig. 2I and fig. S2H; VAm, 15.3%; MDc, 12.5%). Linear trend analysis leveraging a General Linear Model (GLM) and linear contrasts revealed that VA cells achieved selectivity at the shortest latency (100 ms; p<0.001; t>3.4), followed by MD (150 ms; p<0.008; t>3.1), and then pPFC (250 ms; p<0.01; t>2.8). The overall latency followed the pattern: VA<MD<pPFC (p<0.032; t>2.4; Fig. 2J). Second, using an abstract rule-selectivity index (SI) ^36,37^, we identified single neurons in each ROI as abstract rule-selective, and calculated the population average abstract rule SI timecourse for each ROI (Fig. 2, D and H). Results from both methods suggested that MDc and VAm populations carry just as much, if not more rule information compared to PFC populations.

**Figure 2.**
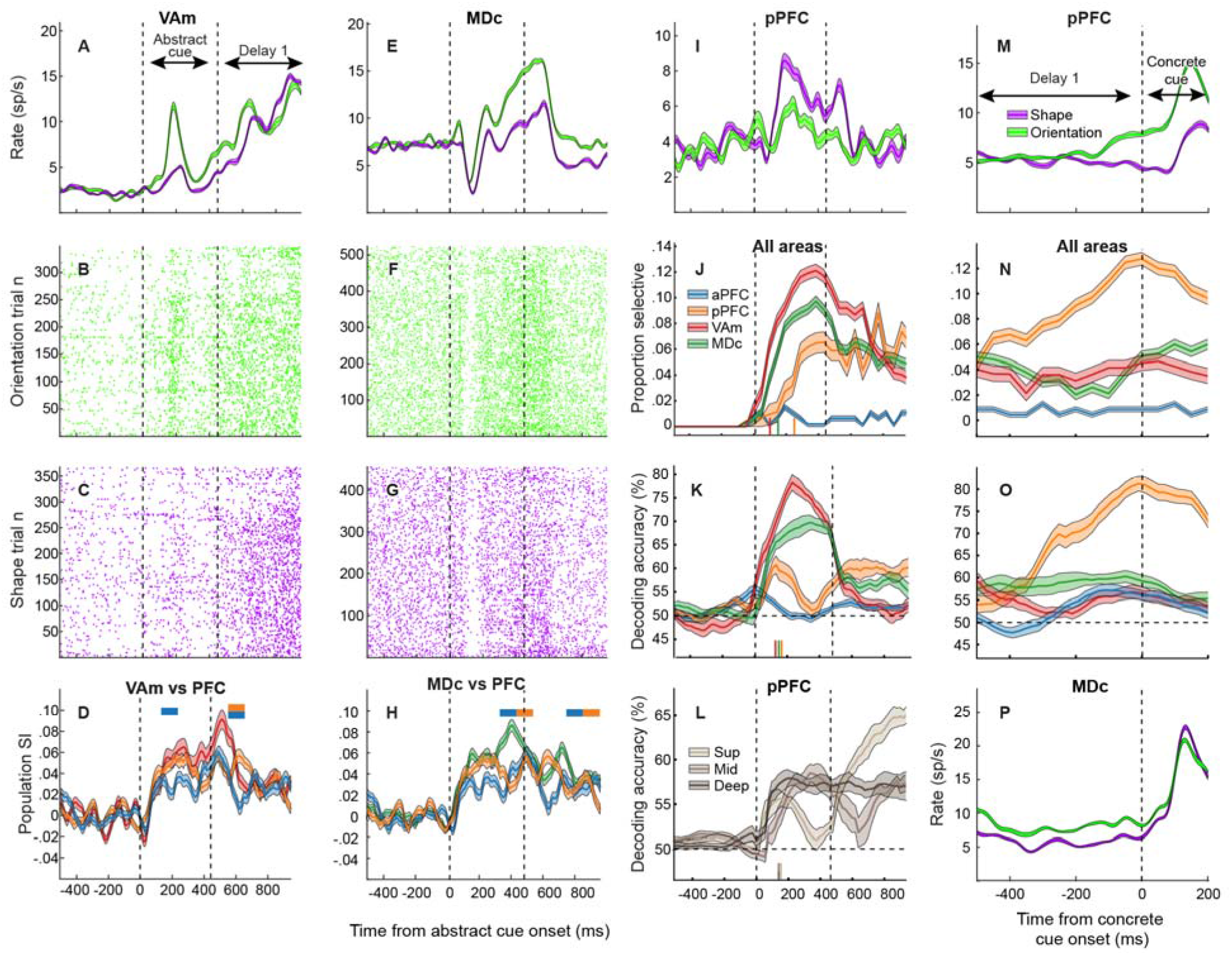
Abstract rule-selectivity emerges first in thalamus during correctly performed trials. (**A** to **L**) Data and results aligned to onset of abstract rule cue and beginning of delay 1; (**M** to **P**) data and results aligned to onset of concrete rule cue (end of delay 1). Single neuron spike density functions (SDFs), raster plots and population SI timecourses for VAm (**A** to **D**) and MDc (**E** to **H**). Color coding legend from panel J is applicable to panels D, H, K, N and O. Colored horizontal bars indicate significantly different 100-ms windows for each PFC region compared to the thalamic nucleus (p < 0.05, one-tailed t-tests). (**I**) pPFC SDF. (**J**) Proportion of rule-selective neurons in each ROI calculated in sliding 100-ms window, aligned to abstract rule cue onset. (**K**,**L**). Timecourses of rule (shape vs orientation) decoding accuracy for each ROI (K) or pPFC layer (L), aligned to abstract rule cue onset (50% accuracy represents chance performance). (**M**) pPFC SDF. (**N**) Proportion of rule-selective neurons in each ROI calculated in sliding 100-ms window, aligned to concrete rule cue onset (end of delay 1). (**O**) Timecourse of decoding accuracy for each ROI, aligned to concrete rule cue onset (end of delay 1). (**P**) MDc SDF. Vertical lines (J-L) above time axes indicate latencies to population abstract rule-selectivity or significant abstract rule decoding.

To further explore this possibility, we decoded the abstract rule from pseudopopulation spiking activity of all neurons for each ROI in all correct trials across all recording sessions (Fig. 2K; fig. S3). Abstract rule information in thalamic populations was greater, and available earlier, than that from PFC populations: During abstract rule instruction (0-450 ms), VAm had the highest decoding accuracy (t > 5.1; p < 0.0007; N=100000; ANOVA) and shortest latency (t > 3.4; p < 0.002; N=100000; ANOVA, 115 ms), followed by MDc (magnitude, t > 3.1; p < 0.006; latency, t > 2.88; p < 0.01; N=100000; ANOVA, 135 ms) then pPFC (magnitude, t > 6.2; p < 0.0001; latency, 150 ms), while aPFC was not informative. This suggests that PFC neurons inherit robust rule-selectivity from thalamic neurons; with BG input possibly contributing to the prominent early role of VAm. Consistent with this, in the pPFC, abstract rule information first appeared (latency 155 ms) in superficial (t > 1.89; p < 0.02; N= 100000; ANOVA) and deep (t > 2.23; p < 0.02) layers – both targeted by VAmc – followed by middle layers (t > 1.45; p < 0.04; latency 165 ms) – targeted by MDpc (Fig. 2L). Initial poor followed by subsequent strong (shown below) abstract rule decoding from PFC is consistent with early weak representations in PFC which are subsequently strengthened, presumably via thalamic influences, when considering the timecourse of rule-related information available in VAm and MDc.

The number of rule-selective neurons in pPFC increased across the first delay when monkeys had to keep the abstract rule in working memory (Fig. 2, M and N). Decoding the abstract rule during this delay (Fig. 2, K and O) suggests pPFC (t > 2.45; p < 0.006; N=100000; ANOVA) and MDc (t > 1.8; p < 0.02) carried more rule information than aPFC and VAm; with pPFC carrying more rule information than MDc (t > 1.3; p < 0.03). Superficial layers of the pPFC were most informative (t > 4.76; p < 0.0003; Fig. 2L) and MDc maintained rule information across the delay (Fig. 2, O and P). As a control for possible differences in spiking variability between areas which could influence decoding accuracy, we calculated the mean-matched Fano Factor ^38^ and there was no significant difference between the areas (fig. S4). This suggests recurrent PFC-MDc loop activation may help to sustain the abstract rule once it has been established by transient VAm activity.

If the thalamus is important for applying abstract rules, then one might expect abstract rule information in the thalamus to be reduced in error trials. Indeed, during abstract rule instruction (0-450 ms), the proportion of rule-selective neurons dropped markedly in error trials (aPFC, 1.4%; pPFC, 2.3%; VAm, 2.1%; MDc, 2.2%; Fig. 3A cf. Fig. 2J). When we trained and tested a decoding model using only error trials, the large peak in VAm and MDc decoding accuracy during correct trials (Fig. 2K) disappeared in error trials (correct > error: VAm, t > 42.78, p < 2.3×10^−16^; MDc, t > 37.29, p < 4.4×10^−13^; Fig. 3C). Consequently, if abstract rule representations in PFC depends on the thalamus, then one might expect abstract rule information in pPFC to be reduced in error trials as well, which is what we observed (correct > error decoding: pPFC, t > 4.9, p < 0.01). Notably, during the first delay when the abstract rule must be maintained in working memory, the proportion of rule-selective neurons was greatly reduced in error trials (Fig. 3B cf. Fig. 2N), and the increase in decoding accuracy in pPFC and MDc during correct trials (Fig. 2O) was absent on error trials (correct > error: pPFC, t > 28.43, p < 7.4×10^−11^; MDc, t > 7.3, p < 0.003; Fig. 3D). We also used error trials to test the decoding model trained with correct trials, to determine how often the irrelevant abstract rule was represented in error trials. The irrelevant abstract rule (i.e., “other” rule) was represented more frequently than the currently relevant abstract rule in error trials (Z = 3.65, p = 0.022, N = 10000; Wilcoxon rank sum; Fig. 3E). Taken together, these results suggest that PFC representations of the abstract rule rely on the thalamus first identifying the relevant rule and then helping to maintain it.

**Figure 3.**
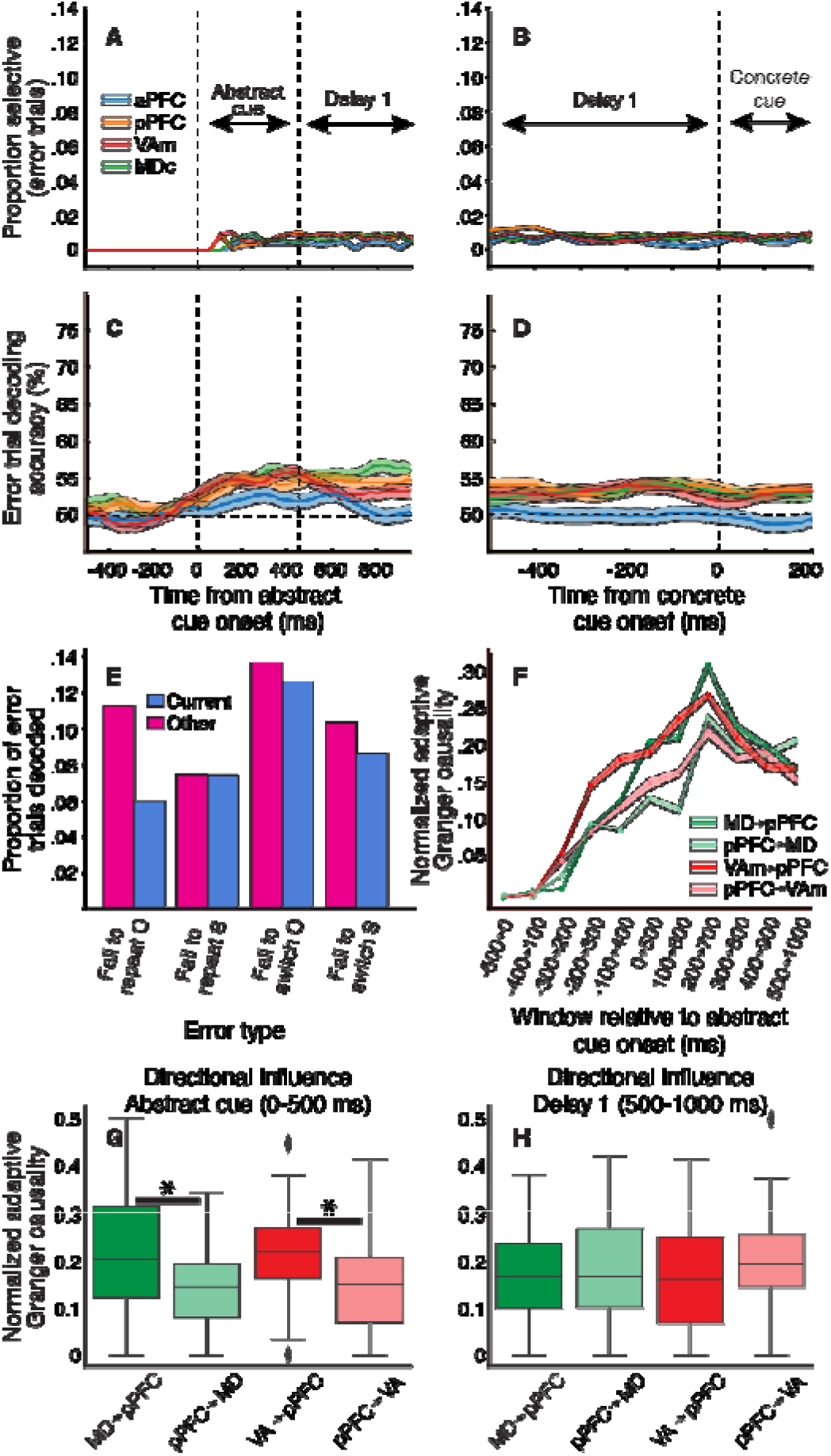
Thalamus causally influences PFC during correct abstract rule processing; but rule information perturbed on error trials. (**A**,**B**) Proportion of rule-selective neurons in each ROI (calculated in sliding 100-ms window), during error trials, aligned to abstract rule cue onset (A) and concrete rule cue onset (B). (**C**,**D)** Timecourse of abstract rule decoding accuracy for each ROI on error trials, by decoding model that was trained on error trials. Timecourses are aligned to onset of abstract rule cue (C) and onset of concrete rule cue at the end of delay 1 (D). Color-coding legend from A also applies to other panels. 50% decoding accuracy represents chance performance. Abstract rule information reduced on error trials (cf. model trained and tested with correct trials in fig. 2, K and O). (**E**) Abstract rule decoded in corticothalamic circuits (mean of all areas) during different types of error trials. Decoding of shape (S) vs orientation (O) in error trials by model trained on correct trials (i.e., model used in fig. 2, K and O). Decoded rule either current rule (i.e., correct rule on current trial; purple) or other rule (rule monkeys erroneously applied, inferred by their executed choice; pink). Behavioral error types based on the correct/rewarded rule (S or O) on previous trial: failed to switch or repeat. (**F**) Mean population timecourse of directed influences between PFC and thalamus aligned to abstract rule cue onset, measured using adaptive Granger causality (AGC) in sliding 500-ms window. (**G**,**H**) Distribution of AGC values between PFC and thalamus during abstract rule cue presentation (G) and delay 1 (H). Box plots display the median and the interquartile range (IQR), which spans from the 25th percentile (first quartile) to the 75th percentile (third quartile). The whiskers extend to the smallest and largest values within 1.5 times the IQR from the quartiles, indicating variability outside the interquartile range. Data points beyond the whiskers, shown as diamonds, represent outliers. Bars with asterisks indicate significant differences between thalamocortical and corticothalamic directional influences (p < 0.05).

To further characterize thalamic influence on abstract rule processing in thalamo-PFC networks, we calculated the adaptive Granger causality between thalamic and PFC neurons, based on their spiking activities ^39^. Across all sessions, causal influences increased between thalamus and cortex with abstract rule cue presentation, peaking around the start of the delay period, and remaining elevated thereafter (Fig. 3F). These results also suggested that directional influences in the thalamocortical direction exceed those in the corticothalamic direction shortly after abstract rule presentation. During abstract rule instruction (0-500 ms), VAm and MDc had significant influence on pPFC (greater than chance and the 500 ms baseline period prior to abstract cue onset; FDR selection of influences with p < 0.05; Fig. 3G). Moreover, both VAm (t=5.18; p=0.001; N=13 sessions; 2073 cell pairs) and MDc (t=4.61; p=0.004; N=22 sessions; 4564 cell pairs) had greater influence on pPFC than the pPFC had on VAm or MDc (which was also above chance and the baseline period). During the subsequent delay period (500-1,000 ms after abstract rule cue onset), MDc influence on pPFC and pPFC influence on MDc were similar (and VAm influence on pPFC reduced, in comparison to the cue period; Fig. 3H). Overall, this supports thalamic causal influence on PFC during abstract rule identification (Fig. 3G), and reciprocal influences between MDc and pPFC during rule maintenance (Fig. 3H).

### Concrete rule-selectivity first appears in PFC

Both PFC and thalamic neurons exhibited robust, short-latency concrete rule-selectivity (Fig. 4, A-D, and fig. S5). The nonparametric tests indicated neurons in the pPFC (23.6%), VAm (8.2%) and MDc (12.2%) were selective for the concrete rule. This concrete rule-selectivity could arise first in the thalamus via convergent cortical inputs (similar to what we observed for the abstract rule), or it could arise first in PFC, if the rule set bias is already established in the PFC following abstract rule processing. The latter would enable sensory inputs to effectively trigger PFC representation of the relevant concrete rule. We performed a population analysis of concrete rule SI in each ROI, to test whether concrete rule information in PFC and thalamus differs (Fig. 4, E and F). This suggested that pPFC neurons may carry more concrete rule information than those of VAm and MDc (Fig. 4E and F); during cue presentation, MDc may carry more information than aPFC (Fig. 4F).

**Figure 4.**
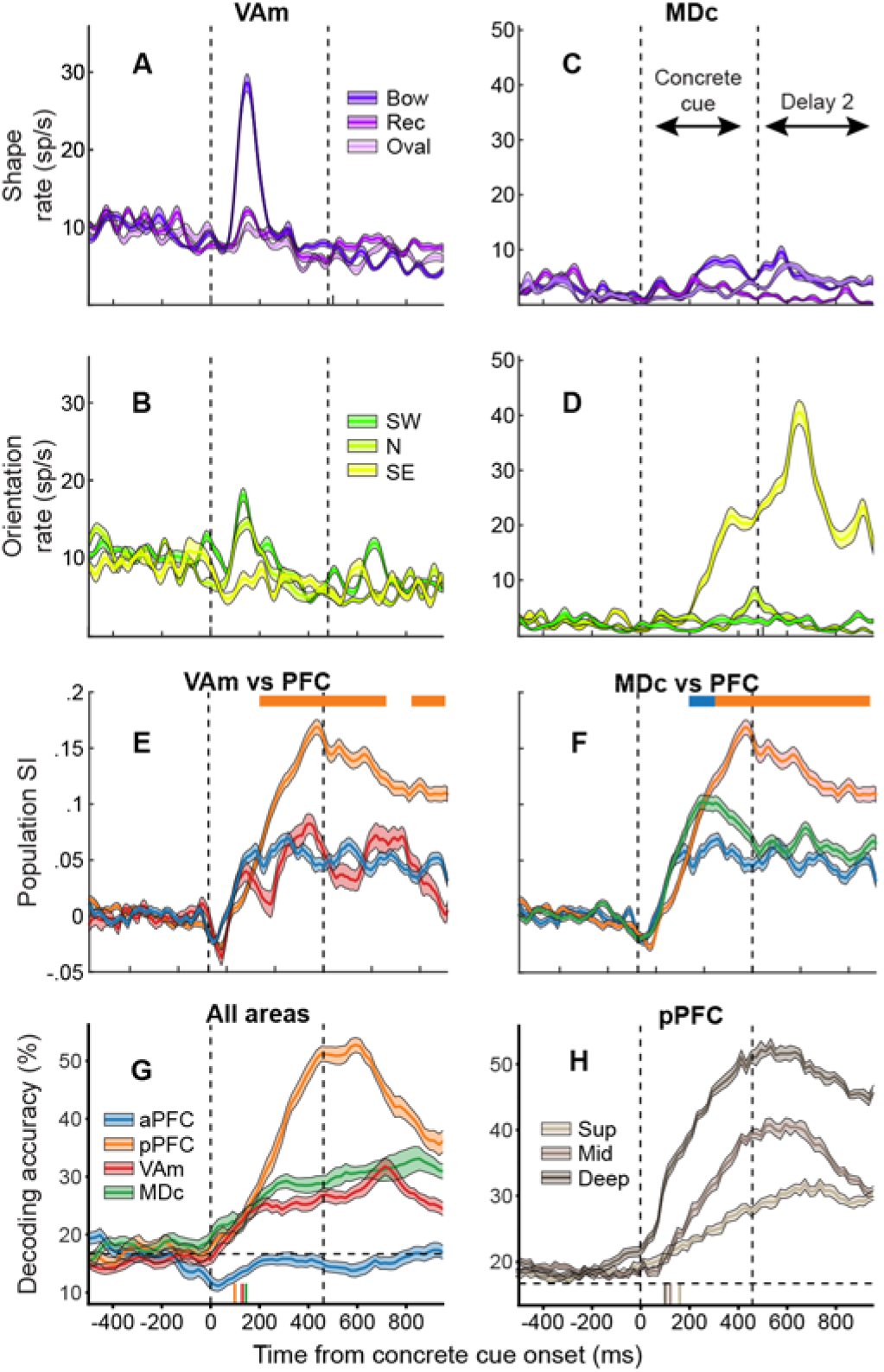
Concrete rule-selectivity arises first in pPFC. SDFs for single VAm (**A** and **B**) and MDc (**C** and **D**) neurons. Population SI timecourses for VAm (**E**) and MDc (**F**), compared with those from PFC. Color-coding legend from panel G also applies to panels E and F. Colored horizontal bars indicate significantly different 100-ms windows for aPFC (p < 0.02, 2-tailed t-tests) and pPFC (p < 0.03, 2-tailed t-tests) compared to the thalamic nucleus. Timecourses of rule (bow, rec, oval, SW, N or SE) decoding accuracy for each ROI (**G**) or pPFC layer (**H**). 16.7% accuracy represents chance performance. Vertical lines above time axes (G,H) indicate latencies to significant concrete rule decoding.

The pseudopopulation decoding analysis confirmed that the pPFC population carried concrete rule information earliest (t > 3.15; p < 0.004; N=100000; ANOVA; latency 105 ms) and most robustly (t > 6.2; p < 0.00005; N=100000; ANOVA) during both concrete rule cue presentation and the second working memory delay period (Fig. 4G). MDc was next most informative (t > 4.87; p < 0.0003; latency, 145 ms) followed by VAm (t > 3.02; p < 0.008; latency, 130 ms), and aPFC was not informative about the concrete rule. Confusion matrices for each ROI demonstrated decoding of each of the six concrete rules in thalamic ROIs and pPFC (Fig. S6). In pPFC (Fig. 4H), concrete rule-selectivity appeared first (t > 5.78; p < 0.00004; N=100000; ANOVA; latency, 105 ms) and most robustly (t > 3.008; p < 0.0008; N=100000; ANOVA) in the deep layers, followed by middle layers (latency, 125 ms), then superficial layers (latency, 160 ms). This is consistent with an intracortical process first giving rise to PFC concrete rule-selectivity, after which core thalamic neurons (particularly in MDc) projecting to PFC middle layers help to sustain concrete rule representations. Thalamic matrix cells projecting to PFC superficial layers seem to play a less prominent role in generating concrete (as compared to abstract) rule-selectivity.

### Strong saccade/target directional selectivity in pPFC supported by MD

Previous work has characterized directional information about visual stimuli/targets and saccadic eye movements in spiking activity of both lateral PFC and thalamus ^17^. Consistent with this precedent, here, single neurons in each ROI also exhibited direction-selectivity based on nonparametric tests: aPFC, pre 4.8%, peri 5.4%, post 7.9%; pPFC, pre 30%, peri 18.7%, post 22.9%; VAm, pre 15.9%, peri 14.7%, post 11.6%; MDc, pre 20.4%, peri 15.7%, post 11.3% (Fig. 5, A to C, and fig. S7). Further, we calculated the directional SI timecourse for single neurons, and for those found to be direction-selective using this method (during at least one of pre-, peri- and post-saccade windows), we calculated a population direction SI. This suggested that pPFC carried more directional information than both thalamic nuclei prior to, during and following saccades (Fig. 5, D and E, p < 0.01, one-tailed t-tests). Both thalamic nuclei carried more directional information than aPFC during at least part of the pre-saccade window (Fig. 5D and E, p < 0.05, two-tailed t-tests). Qualitatively speaking, according to this directional SI, during the peri-saccade window, directional information appeared stable across all ROIs except VAm, whose directional SI population curve dips below that of aPFC (p < 0.05, two-tailed t-test). After saccade onset, pPFC directional information appears to peak, and then to decrease along with that expressed in MDc, while directional information was similar for aPFC and VAm.

**Figure 5.**
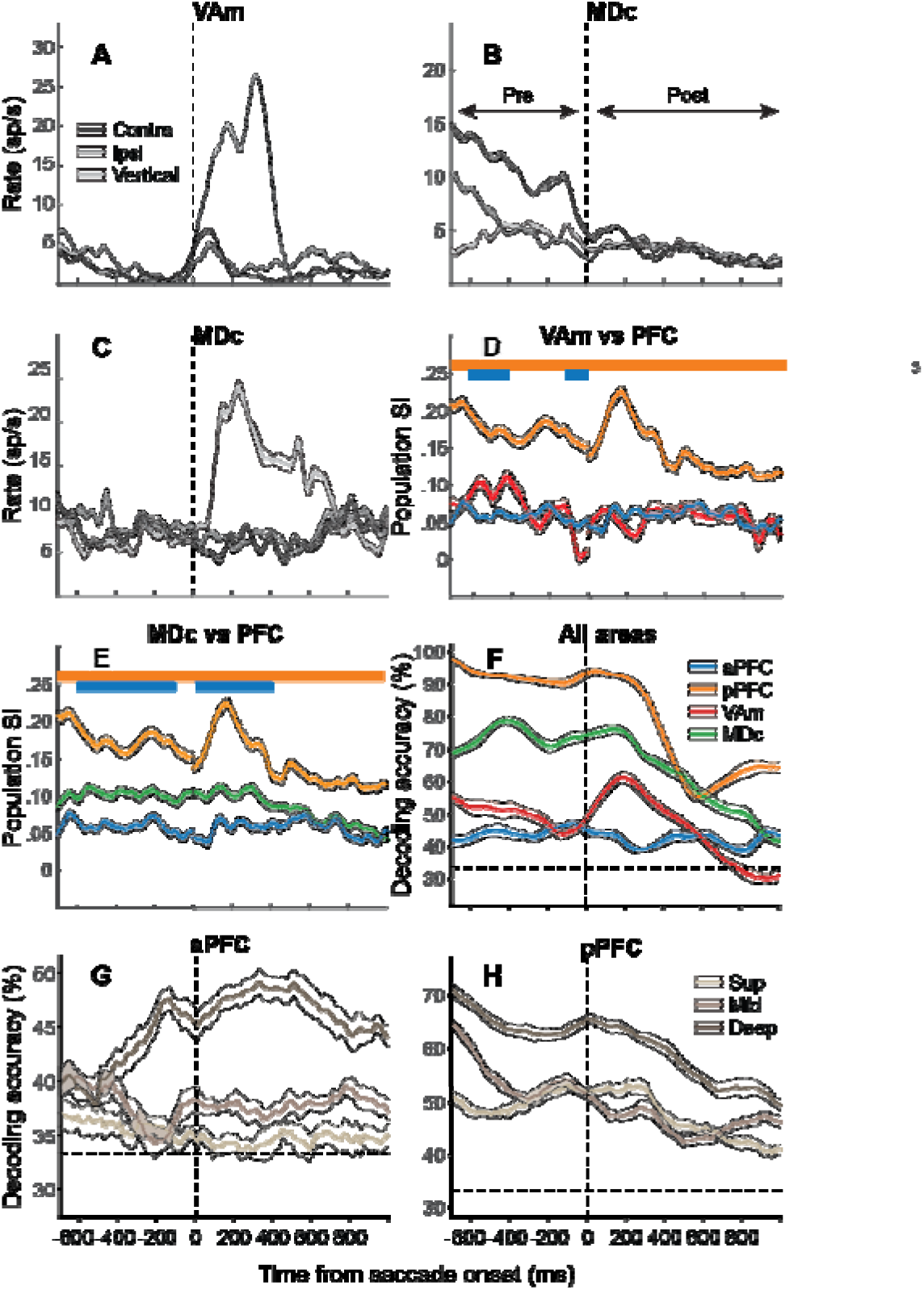
Target/saccade directional selectivity strongest in pPFC followed by MD. (**A**-**C**) SDFs for VAm (A) and MDc (B and C) neurons aligned to saccade onset. (**D**,**E**) Population SI timecourses for Vam (D) and MDc (E), compared with those from PFC. Colored horizontal bars indicate significantly different 100-ms windows for aPFC (p < 0.05, two-tailed t-tests) and pPFC (p < 0.01, two-tailed t-tests). (**F**-**H**) Timecourses of direction (left, up or right) decoding accuracy for each ROI (F) or PFC layer (G and H). 33.3% accuracy represents chance performance.

The pseudopopulation decoding analysis of directional-selectivity aligned with our univariate results. pPFC carried the most robust directional information followed by MDc, VAm then aPFC (t > 2.99; p < 0.004; N=100000; ANOVA; Fig. 5F). Confusion matrices for each ROI confirmed decoding of each direction (fig. S8). The robust directional information in pPFC reflects greatest information carried by its deep layers, followed by middle then superficial layers (t > 3.63; p < 0.001; N=100000; ANOVA; Fig. 5H). Similarly, in aPFC, directional information was greatest in deep, then middle, and then superficial layers (t > 3.4; p < 0.002; N=100000; ANOVA, Fig. 5G). These patterns are consistent with PFC output from deep layers driving directional (choice) information in their target regions before, during and following saccades.

### Outcome information processed in MD-PFC circuits

One important aspect of cognitive control is monitoring whether chosen actions achieve desired outcomes ^40,41^. Many neurons in each ROI had activity modulated by trial outcome (Fig. 6, A to F, and fig. S9). Outcome-selective neurons were detected in every ROI based on nonparametric tests (aPFC, 40.1%; pPFC, 25.4%; VAm, 21.0%; MDc, 52.0%). Some neurons showed greater activity following errors compared to correct trials (Fig. 6A, fig. S9C, F), while others showed the opposite pattern (Fig. 6D, fig. S9B, E), and yet others showed a more complex dynamic modulation pattern (fig. S9A, D). Some neurons showed sustained modulation by trial outcome, persisting across the intertrial interval (Fig. 6D, fig. S9C). Such a pattern may indicate an influence on the allocation of cognitive resources on the following trial or updating of reward expectations to optimize decision-making.

**Figure 6.**
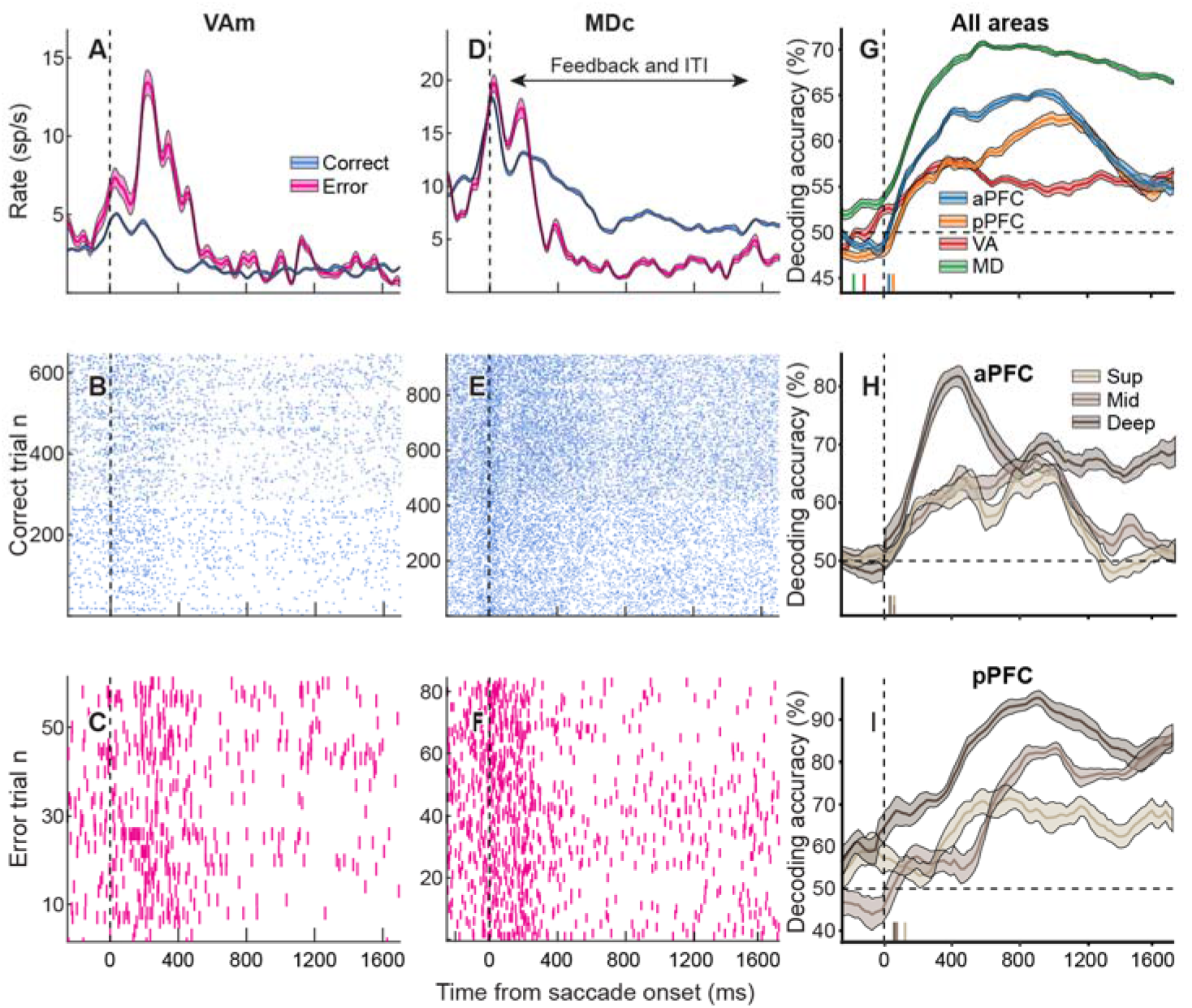
Outcome-selectivity arises first in thalamus. (**A**-**F**) SDF and raster plots for single VAm (A to C) and MDc (D to F) neurons aligned to saccade onset. (**G**-**I**) Timecourses of outcome (correct vs error) decoding accuracy for each ROI (**G**) or PFC layer (**H** and **I**). 50% accuracy represents chance performance. Vertical lines (G-I) above time axes indicate latencies to significant outcome decoding.

The pseudopopulation decoding analysis showed that each ROI carried information on trial outcome, with MDc expressing this information earliest, followed by VAm, aPFC, then pPFC (t > 0.67; p < 0.03; N=100000; ANOVA; Fig. 6G). During the first 1000 ms post-saccade, MDc carried the most information on outcome, followed by aPFC, pPFC, then VAm (t > 1.9; p < 0.003; N=100000; ANOVA). Beyond this temporal window, MDc continued to carry the most information, while aPFC, pPFC and VAm populations carried similar outcome information (t > 2.93; p < 0.0001; N=100000; ANOVA). However, there were also differences in decoding outcome by PFC layer (Fig. 6, H and I).

In aPFC and pPFC, outcome information emerged earliest (t > 1.32; p < 0.01 and t > 1.2; p < 0.03, respectively) and was strongest (t > 2.1; p < 0.007 and t > 2.54; p < 0.005, respectively) in deep layers (Fig. 6H, I), with decoding accuracy comparable to that measured from MDc (Fig. 6G). In pPFC, middle layers carried greater outcome information than superficial layers (t > 2.54; p < 0.005; Fig. 6I). This is consistent with recurrent thalamo-PFC loop activation involving thalamic core neurons sustaining outcome information. The latency results suggest that this information may originate in MDpc ^42^.

### PFC-BG-thalamus model reproduces *in vivo* activity

To characterize the mechanism by which selection and maintenance of abstract rules occurs, we built a layer-specific PFC-BG-thalamic model comprising leaky integrate-and-fire neurons, to process the abstract rule cue and working memory information across the subsequent delay period in the HRT (Fig. 7A, table S1 and table S2). We decoded the abstract rule from the spiking activity of modeled neurons for each ROI. Initially after cue onset, superficial layer PFC neurons showed sensory-selective responses (i.e., raster responses to one of the four cues only), with inhibition during non-preferred cue presentation (fig. S10, A to D). Deep layer PFC neurons showed relatively less sensory-selectivity, due to the convergence across PFC layers (fig. S10, E to H and fig. S11). However, abstract rule-selectivity first emerged in the thalamus, enacted by prominent cortico-thalamic convergence (Fig. 7B, and fig. S10, K to N). VAmc had the shortest latency to rule-selective responses (50 ms; t=4.2, p=0.0006), followed by MDpc (150 ms; t=3.81, p=0.001). The cue-induced BG-mediated disinhibition of VAmc produced a facilitating time window which permitted VAmc neuron firing (fig. S10, K and M). Abstract rule information was then sent to PFC (200 ms latency; t=3.22, p=0.002), and maintained across the delay period in PFC-MDpc loops (fig. S10, I, J, L and N).

**Figure 7.**
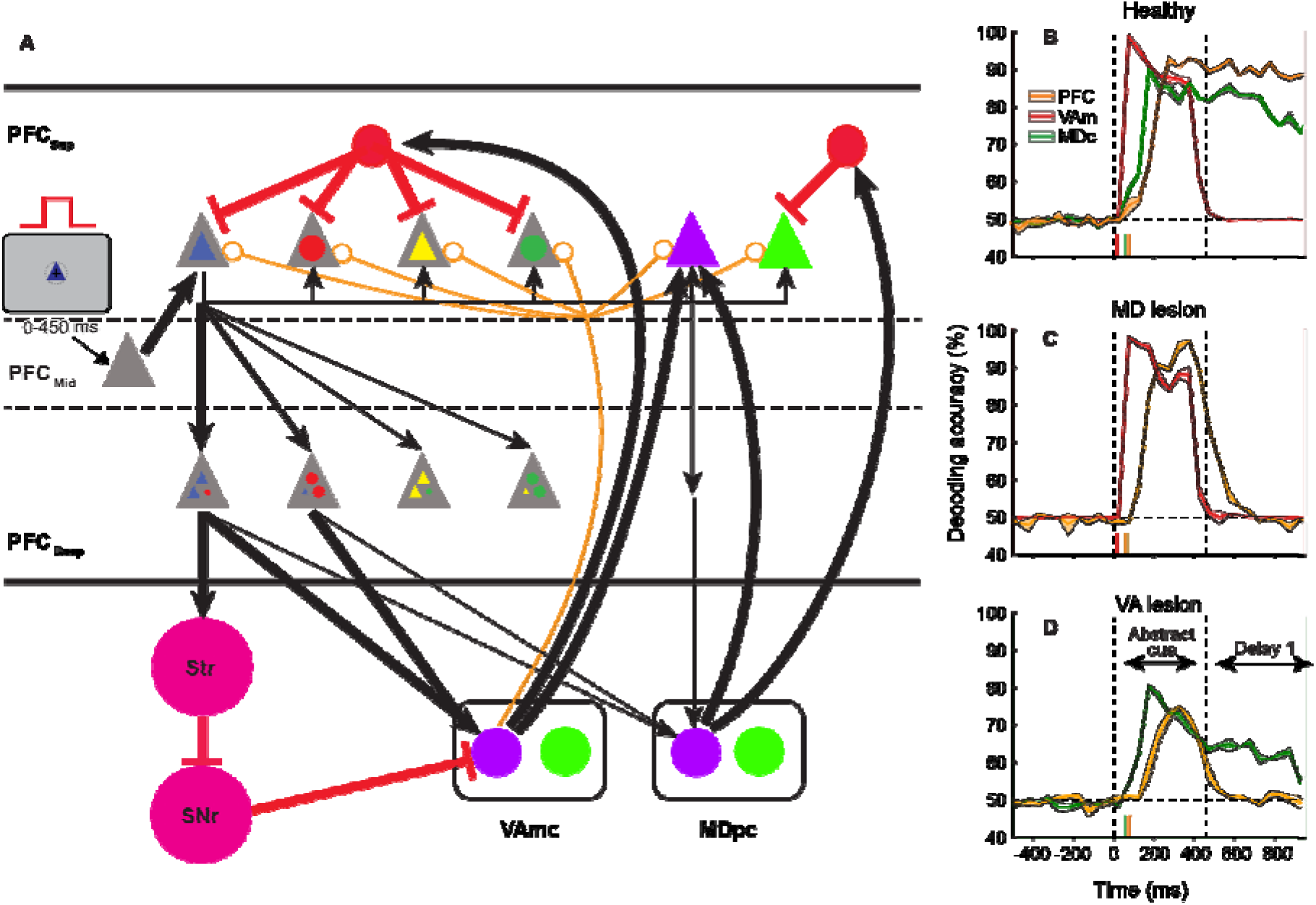
PFC-BG-thalamic model reproduced *in vivo* abstract rule-selectivity and thalamic-lesion modeling disrupted PFC rule representations. (**A**) *in silico* PFC includes superficial, middle (rule cue input layer) and deep layers; BG includes striatum and SNr. VAmc receives disinhibitory input from BG and driving input from PFC deep layers; MDpc receives weaker (reflecting modulatory) input from PFC deep layers. VAmc provides divergent (matrix) and driving (core) input to PFC, and MDpc provides driving input to PFC, reflecting known primate anatomy and physiology ^8^. Flow of information through PFC layers, striatum, SNr and thalamus simulated using leaky integrate-and-fire cells. Each triangle or circle represents a different cell population. PFC superficial layer contains populations selective for one of the four cues (blue triangle, red circle, yellow triangle, dark green circle) or one of the abstract rules (purple, shape rule; green, orientation rule). A population of inhibitory interneurons, driven by VAmc, imposed general inhibition on cue-selective PFC superficial cells; and a population, driven by MDpc, specifically inhibited superficial cells selective for the irrelevant rule. PFC deep layer populations receive driving input from superficial layer cells. VAmc and MDpc cell populations shown are selective for shape (purple) or orientation (green) rules. Black arrows represent excitatory projections, red lines with flat terminations indicate inhibitory projections, and orange lines with open circle terminations represent divergent matrix projections (model connectivity in table S1). Thicker lines represent projections having synaptic weight > 0.003; thinner lines represent projections having synaptic weight ≤ 0.003 (model synaptic weights in table S2). Time courses of abstract rule decoding accuracy from each ROI’s spike data generated by complete, “healthy” (**B**), MD-lesioned (**C**) or VA-lesioned (**D**) models. Color-coding legend in B also applicable to C and D.

### Thalamic lesion modeling disrupted PFC rule representations

We next modeled lesions of VAmc or MDpc by disconnecting them from the rest of the network, to test influences on abstract rule processing in other circuit nodes (Fig. 7, C and D). Modeled VAmc lesions markedly reduced rule-selectivity at all time points after cue onset (although latencies did not differ: MDpc 150 ms; t=2.22, p=0.002; PFC 200 ms; t=2.88, p=0.005). Decoding accuracy dropped about 20% during cue presentation for both MDpc and PFC and, during the delay period, decoding declined further for MDpc and was at chance levels for PFC (Fig. 7D). In comparison, modeled MDpc lesions did not affect rule representations in PFC and VAmc during cue presentation (VAmc 50 ms; t=4.2, p=0.0006; PFC 150 ms; t=3.65, p=0.001). However, decoding was at chance levels for VAmc and PFC during the delay period (Fig. 7C). This is consistent with a mechanism where VAmc driving projections transmit abstract rule information to PFC; and VAmc’s modulatory projections regulate cortico-cortical connections, allowing recruitment of additional PFC neurons to the rule representing ensemble, necessary to bridge the cue and delay periods. MDpc driving input to PFC yields synchronized activation of PFC neurons, which recruits more MDpc neurons to support rule information maintenance across the delay.

### Model MD provides “sweet spot” for cortical excitability

For comparison with the original “healthy” model (our baseline regime), we created model regimes with either relatively weaker weights (more subtle manipulation than lesions) or stronger weights (taking the model in the opposite direction of the lesions) for the MDpc projection to PFC rule-selective neurons. For weaker MD-PFC weights, there was still PFC rule-selectivity during the abstract rule instruction, but not during the subsequent delay period (fig. S12). For stronger MD-PFC weights, we found increased PFC activity for all previously rule-selective neurons, irrespective of the relevant abstract rule. This reduced rule-selectivity during trials, and gave rise to inappropriate activations outside trials (fig. S12). In the weak or strong regime, the signal-to-noise ratio was reduced (cf. our baseline regime), either due to reduced rule signal or increased noise respectively. Consequently, working memory activity in PFC was perturbed. This and the model lesions suggest that in model regimes with moderate thalamo-cortical weights (our baseline regime), both VAmc and MDpc were needed to select and maintain abstract rules, highlighting the importance of matrix-like neurons to promote and shape PFC processing modes, in addition to core-like neurons supporting ongoing operations.

### Error trials due to perturbed rule and delay thalamo-cortical activity in model and *in vivo*

To investigate possible mechanisms underlying behavioral errors in macaques, we expanded the healthy model to additionally process the concrete rule and output the direction (contra, ipsi, vertical) of the choice. In the model (fig. S13), the abstract rule-selective PFC neurons drive inhibition of the irrelevant concrete rule-selective PFC neurons, typically resulting in the concrete cue activating the relevant concrete rule-selective PFC neurons and their associated choice-selective neurons. The model determined the choice based on the maximally activated group of direction-selective PFC neurons. In a small subset of trials, the model output the incorrect choice, akin to behavioral errors in macaques. The main error type (4.6% of all simulated trials) was due to a breakdown in abstract rule processing during the first delay period (fig. S14, left columns under “failure of delay maintenance”). This occurred when there was initial activation of both shape and orientation abstract rule-encoding neurons (when SNr-mediated inhibition was reduced), which led to mutual inhibition of PFC ensembles representing each abstract rule. This is consistent with the reduced abstract rule-selectivity and reduced delay activity *in vivo* (Fig. 3A-D and fig. S14, bottom row). More rarely (0.2% of all simulated trials), the second error type was due to representation of the incorrect abstract rule (fig. S14, right columns under “elevated activity for wrong rule”). This occurred when there was increased (stochastic) activity of abstract rule-encoding thalamic neurons selective for the incorrect or other rule, just prior to, and at onset of, the abstract rule cue. This is consistent with the increased activity of a subset of abstract rule-selective neurons for their non-preferred rule during error trials *in vivo* (Fig. 3E and fig. S14, bottom row). Taken together, the model and *in vivo* results suggest that perturbed thalamo-cortical activity can give rise to behavioral errors.

## DISCUSSION

Overall, we have shown that robust abstract rule information is first available in the thalamus, especially matrix-heavy VAm, which is then issued to PFC superficial layers, to establish a cortical bias for the relevant rule set. The connectivity patterns between PFC and thalamus feature convergent patterns of cortical inputs to thalamus, which enable abstract rule selection, and divergent thalamic outputs to PFC. which disseminate the rule ^8^. Once this occurs, circuits involving MD core and PFC neurons may provide the drive necessary to maintain the bias for the relevant abstract rule, facilitating optimal future computations. Specifically, this bias may poise the system to efficiently transform subsequent inputs to subordinate rules and responses, once cognitive control demands lessen and more concrete, stimulus-mapped behavior is sufficient. Our model suggests that the thalamus ensures a “sweet spot” for cortical excitability and connectivity, in which cortical neurons can be flexibly recruited in different tasks and, in each task, be noise-robust, maintaining signal fidelity.

Such a cortico-thalamo-cortical architecture offers advantages over a purely cortico-cortical architecture. The far greater number of neurons in cortex enable a large repertoire of detailed representations, while the signal compression via converging cortico-thalamic inputs ^43^ enables extraction of latent variables (in a sense similar to this operation in an autoencoder), or rules, in the thalamus. Subsequently, the thalamus can set up task sets across cortex based on the relevant rules, via the extensive divergent thalamo-cortical connectivity. Thus, the cortex and thalamus together can give rise to flexible implementation of a broad array of behaviors in response to an endless assortment of possible situations.

Previous work demonstrated that activity in superficial PFC layers reflects maintenance of information during working memory ^35^. Similarly, in our study, maintained abstract rule information was greatest in superficial PFC layers. However, when the information held in working memory was more concrete, i.e., closer to specifying an action, deep PFC layers assumed a greater role (cf. superficial layers). Deep PFC layers also had a larger role in representation and maintenance of behavioral outcome information. This suggests that ongoing activity in superficial versus deep layers of PFC preferentially carries different types of information ^44^– sensory/cognitive and action/outcome respectively – reflecting the canonical mechanisms of feedforward sensory input via superficial layers and the descending motor-related output from deep layers ^45^.

It has been reported that VA neurons encode stable stimulus values ^46^, and that MD neural activity is predictive of behavioral outcome ^47^. Here we show that thalamus represents behavioral outcome information early, and before aPFC, suggesting thalamus may inform cortex about variables linked to the need for adjustments in allocation of cognitive control, and thus prompts a necessary revision to current frameworks emphasizing orbital and medial frontal cortical contributions to these processes ^8^. This aligns with the anatomical work showing primate MD, along with PFC deep layers, receives dense innervation from midbrain dopaminergic centers ^19,48^, perhaps associated with unique dopaminergic mechanisms relative to those in BG, which has a well-established role in reward processing and reinforcement learning ^20,21^. This would also be consistent with observations in human patients with thalamic lesions impacting MD and VA, who exhibited atypical performance monitoring-related behavior and event-related potentials ^22^. Further, non-human primate MD and PFC represented the preceding response and its outcome, needed for updating reward expectations to optimize decision-making and allocation of cognitive resources ^49,50^.

Based on influential models positing that the anterior-to-posterior dimension of PFC is functionally organized according to abstraction level ^2,51^, we expected aPFC to strongly encode abstract rules, relative to pPFC. In contrast, we found aPFC carries little information about specific task events, and instead appears to play a prominent role in monitoring outcomes, consistent with aPFC supporting metacognition ^52^. pPFC, on the other hand, represented both abstract and concrete rules, inconsistent with a PFC topography based on policy abstraction level. However, the aPFC role in outcome monitoring at longer timescales suggests there may be a topography based on temporal abstraction ^51,53^. Moreover, the time-course of outcome information in the cortex is consistent with a spread from anterior-to-posterior PFC, either directly or via thalamus.

Because VAm and MDc selected and maintained abstract rules, models characterizing the functional organization of the PFC need to incorporate thalamic loops. These loops enable sub-cortical areas to select relevant abstract information from high-dimensional cortical representations, and subsequently update PFC representations to shape task sets, thus playing a vital role in cognitive control.

## ACKNOWLEDGMENTS

This work was supported by National Institutes of Health grants R01MH110311, R01NS117901 and P51OD011106, Whitehall Foundation grant 2015-12-71, and Brain and Behavior Research Foundation grant 23017.

## AUTHOR CONTRIBUTIONS

Conceptualization: JP, YS; Investigation: JP, NK, MR, SMoh, AC, YS; Methodology, Software, Formal Analysis: JP, MA, SS, ER, XC, MF, SMof, YS; Supervision, Funding acquisition: YS; Writing – original draft: JP, MA, SMof, YS; Writing – review & editing: JP, MA, NK, MR, SS, ER, XC, MF, SMoh, AC, CM, SMof, YS.

## SUPPLEMENTAL INFORMATION

Figures S1 to S14 and Tables S1 to S3

## STAR METHODS

### KEY RESOURCES TABLE

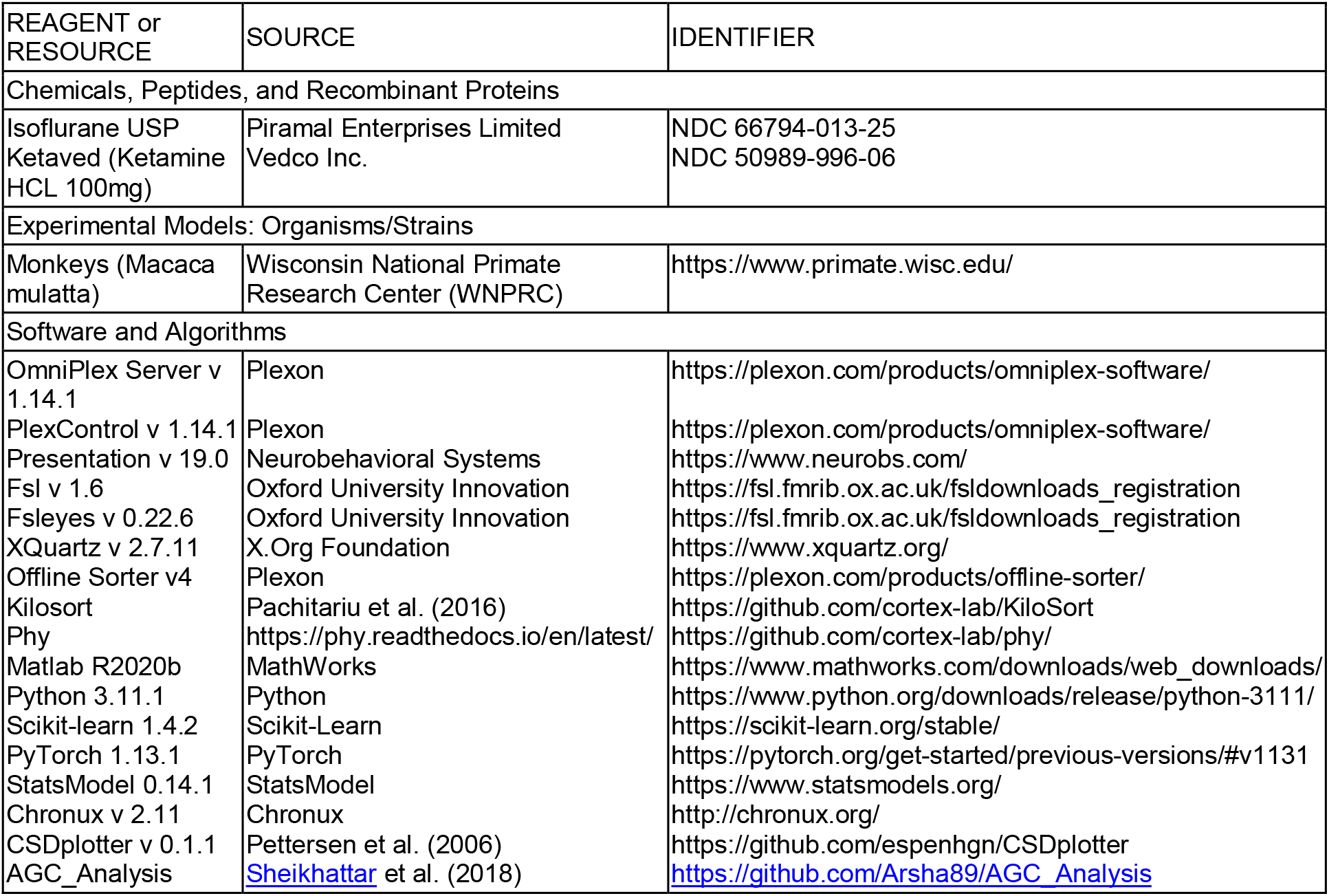

#### LEAD CONTACT AND MATERIALS AVAILABILITY

This study did not generate new unique reagents. Further information and requests regarding resources, equipment, and experimental methods should be directed to, and will be fulfilled by, the Lead Contact, Yuri B. Saalmann.

#### EXPERIMENTAL MODEL AND SUBJECT DETAILS

We acquired data from two male rhesus monkeys (*Macaca mulatta*, age 7 and 9 years, weighing 9.5 and 10 kg, respectively). The daily needs of the animals were met by experimenters and husbandry staff at the Wisconsin National Primate Research Center (WNPRC), where the animals were housed. The health of the animals was monitored by WNPRC veterinary staff. All procedures were approved by the University of Wisconsin-Madison Institutional Animal Care and Use Committee, and conforms to the National Institutes of Health Guide for the Care and Use of Laboratory Animals

### METHOD DETAILS

#### Neuroimaging data acquisition

Prior to implant surgery, we acquired a high-resolution structural brain image to delineate thalamic and cortical regions of interest (ROIs). We also acquired diffusion-weighted MRI data, to identify interconnected network sites and target subregions of interest inside thalamic nuclei. Following craniotomy, we acquired additional structural brain images with electrodes *in situ* to confirm their positioning.

We performed diffusion-weighted and structural imaging on each monkey using the GE MR750 3T scanner (GE Healthcare, Waukesha, WI). Monkeys were pre-medicated with ketamine (up to 20 mg/kg body weight) and atropine sulfate (0.03-0.06 mg/kg) prior to intubation. Subsequently, isoflurane was administered (1-3% on ∼1 L/min O_2_ flow) with a semi-open breathing circuit and spontaneous respiration, to maintain general anesthesia. We monitored the monkeys’ vital signs (expired carbon dioxide, respiration rate, oxygen saturation, pulse rate, temperature) using an MR-compatible pulse oximeter and rectal thermometer.

For the three-dimensional T1-weighted structural volumes, we used an inversion-recovery prepared gradient echo sequence with the following parameters: FOV = 128 mm^2^, matrix = 256 × 256, number of slices = 166; 0.5 mm isotropic; TR = 9.68 ms; TE = 4.192 ms; flip angle = 12°; and inversion time (TI) = 450 ms. We collected 10 T1-weighted structural volumes and calculated the average image for each monkey using the FMRIB Software Library (FSL) ^54^. During diffusion scan sessions, this protocol was used to acquire an additional pair of structural volumes for co-registration purposes. Finally, to localize electrodes, we averaged two structural images with electrodes *in situ*.

We used 2D echo-planar imaging (EPI) with a pulse sequence spin echo gradient pulse ^55^ with parameters: field of view 128 mm x 128 mm, resolution 1.0 mm x 1.0 mm; 80 × 1.0 mm coronal slices; repetition time = 11 000 ms; echo time = 77.5 ms; and parallel imaging (ASSET) factor of 2, to reduce the echo-spacing and thus image distortions. The acquisition used partial Fourier encoding of 0.0625 in the phase encoding direction to reduce the echo time and the resulting images were reconstructed using homodyne processing.

We used two protocols for the acquisition of the diffusion MRI data ^42^. For one monkey, we used protocol A, which involved positioning the monkey in the 16-channel receive-only head coil (MRI instruments) using fitted cushions, and acquiring 8 b=0 images and 60 b=1000 s/mm^2^ images. The entire acquisition was repeated 13 times. We also acquired a B0 fieldmap using a three-dimensional gradient-echo sequence with three echoes and the iterative decomposition of water and fat with echo asymmetry and least-squares estimation (IDEAL) method ^56^, which was used for geometric unwarping of diffusion-weighted data.

For the second monkey, we used protocol B, which involved positioning the monkey in the 8-channel receive-only head coil (Clinical MR Solutions) using a customized MRI-compatible stereotaxic apparatus. The acquisition protocol was similar for this monkey, but with 10 b=0 images and 120 diffusion directions, and with a second set of b=0 images acquired using reversed phase encoding. The entire acquisition was repeated 10 times. The inclusion of the second set of reverse phase encode b=0 images enabled the use of newer utilities for distortion correction (the two acquisition protocols and preprocessing pipelines produced qualitatively similar results, see ^42^).

#### Probabilistic tractography on diffusion-weighted neuroimaging data

We used FSL to preprocess the diffusion MRI data ^57,58^, and to calculate probabilistic connectivity maps using probabilistic tractography. Based on these connectivity maps, we could isolate voxels in the mediodorsal thalamic nucleus (MD) with high probability of connection to the principal sulcus (PS) in the dorsolateral prefrontal cortex (dlPFC), and in VA with high probability of connection to the substantia nigra pars reticulata (SNr), which anatomists have used to define VAmc ^59^. VAmc has preferential connections with the entire PFC ^8,60-62^. Thus, this approach enabled targeting of electrodes to interconnected sites in prefrontal thalamocortical networks based on each monkeys’ specific anatomy.

##### Preprocessing: Correction for motion, eddy currents and susceptibility distortions

For the dataset acquired with protocol A, we used the FSL utility Eddy_Correct to correct for motion (all EPIs) and eddy currents (diffusion weighted). We geometrically unwarped EPIs using the fieldmap and magnitude images acquired in the same session, to correct for susceptibility distortions ^63^. In detail, to obtain a transformation matrix between fieldmap space and diffusion space, we manually skull-stripped the magnitude image. This volume was then forward-warped according to the fieldmap, using FSL’s Utility for Geometrically Unwarping EPIs (FUGUE), and registered (affine with 12 degrees of freedom (DOF) ^64^) to an averaged, skull-stripped non-diffusion reference volume. We then applied the corresponding transformation matrix to the fieldmap image (scaled to rad/s and regularized by a 2-mm 3D Gaussian kernel), aligning it with the non-diffusion reference volume, so that it could be used to unwarp the EPIs with the FUGUE utility. Next, we skull-stripped the T1-weighted structural brain image and co-registered it to the averaged, skull-stripped, corrected non-DWI reference volume (12 DOF) to produce the transformation matrix between the two spaces. The corrected diffusion and b=0 volumes were then averaged across scan repetitions to produce a single set of 68 volumes (8 b=0 and 60 b=1000) for further processing.

For the dataset acquired with protocol B, we used FSL’s Topup utility ^57^ to estimate the field distortions caused by susceptibility artifacts. In detail, we registered all b=0 images acquired with both phase encode directions to create a single pair of images with higher signal to noise ratio for submission to Topup. This pair of distorted images was used to estimate the field, and the two images were combined into a single corrected volume. We then submitted the Topup output, along with the entire concatenated dataset, to FSL’s Eddy utility. Eddy uses the Topup field estimates, and corrects for motion, susceptibility distortions and eddy current distortions simultaneously ^65^. A new set of direction vectors is output by Eddy, which accounts for any motion correction, so these data were not averaged at any stage of processing or analyses. We also co-registered structural space to diffusion space, as described above.

##### Estimation of diffusion parameters and probabilistic tractography

For the probabilistic tractography analyses, cortical and subcortical ROIs were manually delineated in each monkey using the high quality T1-weighted structural brain image and in consultation with stereotaxic atlases ^66-68^. We applied the transformation matrix (from the co-registration of the structural image to the reference non-diffusion image) to the ROI masks for probabilistic diffusion tractography (PDT) analyses.

We performed the PDT analyses using FSL’s Diffusion Toolkit (FDT). The algorithm modeled two fiber populations per voxel ^69^, which is suitable for the complex fiber architecture of the thalamus ^70-72^. The posterior probability distributions of fiber direction at each voxel were calculated, for each monkey ^70,73^. The anatomical plausibility of paths through thalamic and cortical ROIs was assessed as described in ^42^.

To identify MD voxels with a high probability of connection with the dlPFC (area 46, which contains our aPFC and pPFC recording sites), and VA voxels with a high probability of connection with the SNr, we performed a PDT analysis to estimate paths passing through any voxel in the thalamic ROI, and the probability that such paths will pass through a voxel in the non-thalamic ROIs, by running “multiple mask” tractography. Five thousand streamline paths were created from each seed voxel in the thalamic masks by drawing samples from their posterior probability distribution (of directions/angles) associated with the path’s current voxel, and then adding the step to the end of the path based on the drawn angle (0.25 mm step length with maximum of 4000 steps). The sampling was repeated, with paths beginning in seed voxels of the non-thalamic ROI. The proportion of these samples (streamlines) passing through each voxel indicates the probability of connection to it. We used an exclusion mask to isolate ipsilateral connections and to eliminate anatomically implausible paths (see ^42^ for details). We used these paths to isolate “projection zones” in MD (from dlPFC) and VA (from dlPFC and SNr) using the method described in ^42^. The locations of these projection zones were then used to guide planning for the craniotomy surgeries.

#### Surgery

Anesthesia was induced using ketamine (up to 20 mg/kg, i.m.) and general anesthesia maintained with isoflurane (1-2%) during aseptic surgical procedures. We used 12 ceramic skull screws with dental acrylic to affix cranial implants containing a head holder and craniotomy placeholder closed off with a custom plastic recording chamber. The cranial implant enabled behavioral training using video eye tracking. Once monkeys were fully trained on the hierarchical rule task (HRT), a second procedure was performed to place four 2.5 mm craniotomies in frontal and parietal bones. The craniotomies provided access to two cortical ROIs and two thalamic ROIs in the right hemisphere (aPFC, pPFC, MDc and VAm). Coordinates for craniotomies were calculated using the high-quality T1-weighted structural images acquired prior to the surgeries and the position of the projection zones estimated in the PDT analyses of diffusion-weighted data. We fitted each craniotomy with a conical plastic guide tube filled with bone wax, through which linear microelectrode arrays (LMAs) were advanced ^72^. The guide tubes were prefabricated using a model of the skull based on the T1-weighted structural images. We also inserted a titanium skull screw positioned inside the recording chamber to serve as a reference.

#### Behavioral task and stimuli

Our ability to flexibly adapt behavior according to current goals and context is known as cognitive control. It is considered hierarchical, as goals can be selected at different levels of abstraction. An important way to implement cognitive control is to apply rules. To investigate thalamic contributions to hierarchical cognitive control and PFC processing, we designed an hierarchical rule task (HRT). The task required monkeys to selectively process (based on initial abstract rule instruction cue) and map responses from (based on subsequent concrete rule instruction cue) one of two stimulus dimensions (while ignoring the other irrelevant stimulus dimension). Thus, each trial involved the presentation of a colored “abstract rule” cue, which informed the animal about the relevant dimension of the forthcoming white “concrete rule” cue (Fig. 1). A red circle or blue triangle instructed monkeys to follow the abstract shape rule; whereas a green circle or yellow triangle instructed monkeys to follow the abstract orientation rule. The two stimuli to cue each abstract rule served as controls for purely visual responses in neurons. The monkeys had to keep this abstract rule information in mind over a brief delay. Importantly, the abstract rule cue did not specify a behavioral response, but one of two possible rule sets, and was thus a more abstract task parameter than the concrete rule. The concrete rule cue involved presentation of one of three possible shapes (bowtie, rectangle, oval), positioned according to one of three possible orientations (southwest, north, southeast). Each concrete rule cue shape and orientation mapped a response rule onto one of three targets, depending on the currently relevant abstract rule. Thus, monkeys could make their decision and prepare their response once they had been presented with the concrete rule cue. However, they had to maintain fixation over a second variable delay period until the fixation cross was removed and three targets were presented. There were nine possible permutations of the concrete cue (Fig. 1A) and 36 experimental conditions; but at the abstract cognitive level, correct trials could be collapsed into six concrete rule categories (3 shape rules and 3 orientation rules).

The training stages involved focusing on one of the two abstract rule sets, to map a response to either shape or orientation (with the other stimulus dimension fixed, so that they could initially ignore it). The animals learned the first rule set by operant and classical conditioning (fig. S1, top row, “Bowtie only” and “North only”). Initially, there was no memory delay. Once the rule set (e.g., orientation rules) was learned on one stimulus feature (e.g., bowtie), then a second stimulus feature was included in the blocks of trials (e.g., ovals). The monkeys learned the stimulus-response mapping more quickly once they had already learned the rules on one stimulus feature, and thus showed generalization (fig. S1, middle row, “Bowtie + Oval”; “North + Southeast”), a hallmark feature defining abstract rule use. The third stimulus feature (e.g., rectangles) was learned the most rapidly (fig. S1, bottom row). The monkeys were then trained to perform the second abstract rule set (ignoring the first important dimension, e.g., orientation, and now attending to the second dimension, e.g., shape). We observed evidence of behavioral generalization once again (fig. S1). Once both abstract rule sets had been learnt, the monkeys were exposed to both rule sets in the same behavioral training sessions and needed to use the instruction cue to select the abstract rule (which was not necessarily required in prior training sessions). Once the monkeys were able to switch between the two abstract rule sets across blocks within the same session, we then included the two abstract rule sets in the same block, randomly interleaved. The last step in training was to build in the memory delays. Monkeys M and J overall performed at a mean accuracy of 91% and 67% respectively (33% being chance performance) for our physiological recordings. There was no difference between behavioral performance in early versus late recording sessions (rank sum test: Monkey M, p=0.1268; Monkey J, p=0.4894).

During each session, monkeys sat upright in a primate chair (Crist Instruments, Hagerstown MD) with their heads immobilized using a head post/head holder system (Neuronitek, London, Canada) and their eyes positioned 57 cm from a 19 inch CRT monitor (100 Hz refresh rate). The monkeys performed the HRT to earn juice rewards, indicating their target choices using eye movements. Eye position was monitored using a video eye tracker (SMI, Berlin, Germany). Behavioral task events, monitoring of eye position and feedback delivery were controlled by custom-written software in Presentation (Neurobehavioral Systems, Albany CA). Rewards were delivered via a TTL pulse issued to an infusion pump system (Harvard Apparatus, Holliston MA). The PC running Presentation issued TTL pulses to the PC running the neural data acquisition system (Plexon, Dallas TX), to store behavioral and task event information, together with neural data. Recording sessions were typically approximately 2 hours in duration and monkeys performed between 351-1047 trials during a recording session (736 trials on average).

Trials were offered to the monkeys via presentation of a white fixation cross (1.72° x 1.72°) at the center of the screen on an otherwise gray background. Monkeys initiated a trial by foveating the fixation cross, which was maintained on the screen for the duration of the trial until target presentation, to anchor the gaze across cue presentation and delay periods. Eye position was monitored within a 5° x 5° window, and if the gaze position exited the window prior to target presentation, the trial was aborted. After a variable preparatory period (500-1100 ms), the abstract rule cue was presented (either a 2.6° diameter circle or a triangle of 3.4° maximum width, duration of 450 ms, Fig. 1, A and B). This was followed by a variable (500-900 ms) delay period, where the monkey was required to hold the cue information in working memory. Next, the concrete rule cue was presented (either a rectangle, oval or bowtie which were framed by a rectangle of dimensions 7° x 2.4 (for ovals and rectangles) or 3.1° (for bowties), duration of 480 ms, Fig. 1, A and B), which was followed by a second variable delay period (500-900 ms). The width of the concrete cues was subjected to a small amount of jitter (+/-0, 0.35°). The concrete cue was presented at either 340° (“north”), 60° (“southwest”) or 125° (“southeast”). Finally, three identical white circle targets (1.72° diameter) were presented 12 degrees to the left, directly vertical, and right of the fixation cross. The monkey had to combine the information conveyed by the two rule cues to produce a saccade toward the correct target within 700 ms of targets onset. To obtain a reward, the gaze had to land in the correct target window (6° x 6°, centered on the target) and remain inside for 50 or 300 ms. When these conditions were met, correct feedback was signaled to the animal: juice reward delivered and an auditory tone played over a pair of speakers positioned on either side of the monitor. If the gaze left the fixation window, flew through the target window, or an incorrect target was foveated, then a distinct auditory tone sounded over the speakers and the trial was aborted. The duration of the intertrial interval was 1100-1700 ms.

#### Electrophysiological recording

We used LMAs with either 24 contacts, 12.5 µm contact diameter and 200 µm spacing between contacts (Microprobes for Life Science, Gaithersburg, MD), 32 contacts, 12.5 µm contact diameter and 150 µm spacing (Microprobes for Life Science; Microprobes LMAs MRI-compatible), or 32 contacts, 15 µm contact diameter and 150 µm spacing (Plexon V-Probes). LMAs were platinum/iridium and 0.8-1 MΩ impedance. We recorded electrical brain activity (filtered 0.1-7500 Hz and sampled at 40 kHz) using a preamplifier with a high impedance headstage and an OmniPlex data acquisition system controlled by PlexControl software (Plexon). The system also recorded digitized eye position from the eye tracker and behavioral event information issued from our task control software.

##### Electrode array localization

The T1-weighted structural images with electrodes held *in situ* by the custom guide tubes allowed for confirmation of optimal linear electrode placement following craniotomy surgery (Fig. 1D). Specifically, the electrode is not visible in the images, but a susceptibility “shadow” artifact appears along its length with a width of approximately one voxel (0.5 mm^3^ on either side of the electrode). Probe tracks and target depths were calculated based on these images – the individual monkey’s anatomy – with cross-referencing to stereotaxic atlases. We re-positioned electrodes as necessary and re-acquired T1-weighted structural volumes until electrodes were in their desired locations in the thalamus and cortex. Offline, we registered the images with electrodes *in situ* to the high-quality structural image acquired prior to surgery. These procedures enabled targeting of both banks of the PS in the aPFC and pPFC, the PS-connected MDc (putative MDpc) and SNr-innervated VAm (putative VAmc) (Fig. 1, C and D).

We used the depth measurements from structural images to guide initial positions of the linear electrodes across the two PFC and thalamic recording sites. All contacts spanned approximately 4.6 mm. This enabled coverage across both banks of the PS (thus, acquisition of neural data from both area 46d and 46v) and a majority of the dorsal-to-ventral span of the thalamic nuclei. Online, we adjusted electrode position if necessary to maximize the number of contacts showing single- or multi-unit spiking activity. In the cortex, we used current source density (CSD) and analyses of unipolar local field potentials (LFPs) to assign the electrode contacts to cortical layers. We also performed post-mortem histology to reconstruct electrode tracks in one monkey, to corroborate evidence from MRI and electrophysiology. After fixing the brain in 10% neutral buffered formalin, the right hemisphere was sectioned into approximately 5 mm thick coronal slices, embedded in paraffin, then thinly sectioned (8 µm). In proximity to the ROIs, we stained sections with Hematoxylin and Eosin, and visualized sections under a microscope to confirm electrode tracks through our ROIs.

#### Neural data preprocessing

##### Spike data

We used two methods to isolate spiking activity from the acquired wideband data. (i) Using Plexon Offline Sorter software, we bandpass filtered data (250-5000 Hz) to extract potential spiking activity (Butterworth, order 4, zero-phase filter) for spike sorting. We used principal components analysis to extract features of the spike waveforms and a T-distribution expectation maximization algorithm to identify clusters of spikes with similar features. (ii) Using Kilosort ^74^ running in Matlab, with manual curation steps taken in the Phy GUI (https://github.com/cortex-lab/phy/), we performed data preprocessing, spike clustering and template matching. We confirmed that the two methods isolated the same units by running each on the same recording session. We isolated activity from 1094 neurons (229 in aPFC, 340 in pPFC, 196 in VAm, 329 in MDc) across 51 recording sessions (19 in Monkey J, 32 in Monkey M).

##### Local field potentials

We applied a sixth-order Butterworth zero-phase filter to lowpass filter data to 100 Hz for current source density (CSD) analysis and to 200 Hz for relative power analysis. Next, data were downsampled to 1000 samples/second. We then applied linear detrending. To remove powerline noise, we used either the Chronux function rmlinesc to remove significant sine waves from the LFP data for CSD, or an adaptive filter method ^75^. The method began with the utilization of an adaptive notch filter to determine the primary frequency of the interference. Leveraging this identified frequency, we synthesized harmonics using discrete-time oscillators. Subsequently, we used a modified recursive least squares algorithm to estimate the amplitude and phase of each harmonic. We then subtracted and calculated interference from the original data to obtain a cleaner signal. Finally, LFP data were cut into chunks surrounding trial events of interest. To minimize the impact of motion artifacts, we excluded any trials with LFP amplitude greater than 5 times the standard deviation of the mean.

#### Current source density (CSD)

We designated electrode contacts and any isolated single units to either superficial (layers 1-3), middle (400-450 µm span above bottom of layer 4) or deep (layers 5 and 6) layers ^34^ based on inverse CSD analysis ^76^. For this, we used the CSDplotter toolbox for Matlab for calculating the inverse CSD in response to concrete cues in the HRT (and this was confirmed with response to other task events). Linear electrodes measure the LFP, ϕ, at N different cortical depths/electrode contacts along the z axis with spacing h. The standard CSD, C_st,_ is estimated from the LFPs using the second spatial derivative.

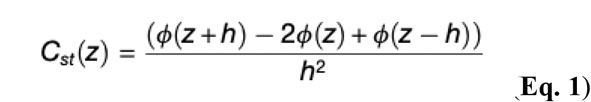

LFPs can also be estimated from CSDs (in matrix form *Φ* = F where Φ is the vector containing the N measurements of ϕ, is the vector containing estimated CSDs, and F is an NxN matrix derived from the electrostatic forward calculation of LFPs from known current sources). The method uses the inverse of F to estimate the CSD, i.e., = *F*^-1^*Φ*. It is assumed that the CSD is stepwise constant between electrode contacts, so the sources are extended cylindrical boxes with radius R and height h.

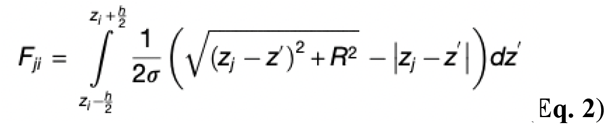

where *σ* is the electrical conductivity tensor and ϕ(z_j_) is the potential measured at position z_j_ at the cylinder center due to a cylindrical current box with CSD, C_i_, around the electrode position z_i_. The inverse CSD provides an estimate of CSD around all N electrode contacts (contrast with standard CSD which does so for N-2) and does not require equidistant contacts like the standard CSD, which permits the analysis of sessions with noisy channels that should be removed.

#### Relative LFP power

We cross-referenced our layer designations estimated using CSD by calculating the relative LFP power across channels/contacts of our cortical LMAs. We analyzed LFP data during the 500 ms prior to abstract cue onset (last 500 ms of preparatory period), which should be weakly stationary. For each channel, we obtained the power spectrum (using the Chronux function mtspectrumc (with time bandwidth product (TW) = 2 and number of tapers (K) = 3) in Matlab), then normalized the power of that channel by that of the channel with the highest power for a given frequency. This revealed the expected common pattern for cortical relative power maps where activity at higher frequencies (50-150 Hz) had higher power in superficial channels than deep ones, while oscillatory activity in the alpha-beta frequencies (10-30 Hz) had higher power in deep channels compared to superficial ones (Fig. 1F), with the characteristic “swoosh” pattern previously described ^23,35^..

#### Selectivity index analyses of spike data

##### Spike rate

We calculated spike density functions (SDFs) to measure the responses of the prefrontal cortical and thalamic neurons. For spike density functions and selectivity index analyses, spikes from each trial were convolved with a Gaussian pulse (λ = 20 ms) and we averaged the resulting functions to produce spike density functions for each neuron, with a temporal resolution of 4 ms.

##### Selectivity index formula

To characterize the response properties of neurons in thalamocortical circuits, we calculated selectivity indices (SIs ^36,37^):

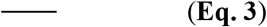

where n is the number of conditions and *λ* is the spike rate in a given condition. We applied this index in a sliding window along 3.0 s chunks of spike data, aligned to various events of interest (window width = 100 ms; step size = 4 ms), thus obtaining a timecourse of SIs from the spike density functions for the abstract rule, concrete rule and direction. To define a selective neuron, we used a threshold which corresponded to baseline mean + 2 standard deviations above this baseline mean, and further the neuron had to exhibit sustained significant SI values for at least 12 consecutive bins (48 ms). We calculated *abstract rule and abstract cue SI* aligned to the abstract rule cue presentation onset, to assess cue-evoked sensory and rule-selectivity as well as rule-selectivity during the first delay period. We calculated *concrete rule and concrete cue SI* aligned to the concrete rule cue presentation onset, to assess cue-evoked sensory- and rule-selectivity. We calculated *directional SI* aligned to the saccade onset to characterize preparatory, peri-saccade and post-saccade directional (saccade or target)-selectivity. Baseline windows were the last 500 ms of the fixation period prior to abstract rule cue onset, for the abstract rule and delay 1, and the last 500 ms of delay 1 for events which followed delay 1, i.e., the concrete rule and target/saccade direction. To define latency to selectivity of a particular single neuron to a visual cue, we found the maximum SI value within the window of interest and calculated the time between event onset and time until the SI reached half of this maximum value. We excluded neurons with unrealistic latencies (i.e., <50 ms).

##### Defining sensory neurons for exclusion from abstract and concrete rule-selectivity index analyses

###### Abstract cues

We included two cues for each abstract rule to enable the dissociation of visual neuronal responses from abstract rule neuronal responses. We ran a sensory SI where the spike rates associated with each of the four abstract cues were input. We reasoned that if a neuron was rule-selective, the cues associated with its maximum two spike rates would reflect the same abstract rule (i.e., red circle and blue triangle, or green circle and yellow triangle). If a neuron was detected as selective based on the sensory SI and failed to meet this rule-related criterion, it was defined as a visual/sensory neuron and excluded from our subsequent abstract rule analyses.

###### Concrete cues

To define sensory responses to the concrete cues, we exploited the fact that (unlike the abstract rule cues), identical concrete rule cues (of which there are 9 permutations) are presented in both shape and orientation trials. We reasoned that if a cell has a maximal response to the same cue under both rule sets, that it is selective for that visual cue rather than any particular concrete rule. In this case, the neuron was excluded from our concrete rule analyses.

##### Population SI timecourse

We calculated the timecourse of SI and assessed for significant SI on individual neurons. If the neuron met criteria for a selective neuron in a particular analysis, we stored its SI data in a population matrix for further processing. For visualization of population SI, we calculated the average SI across each timebin (+/-standard error) and plotted its timecourse. To start probing when selectivity in the thalamus may differ from selectivity in the PFC, we used t-tests to compare selectivity index values obtained from thalamic neurons and PFC neurons in 100 ms-wide non-overlapping jumping windows (p-values were not corrected across these windows).

#### Fano Factor calculation

We employed the methodology delineated by Churchland and colleagues to compute the mean-matched Fano factor for each designated area ^38^. This mean-matched approach controls for any spike rate differences between areas (as Fano factor decreases as spike rate increases). Our timeline of interest was centered around the abstract rule cue, spanning from 100ms before to 1000ms after the cue’s onset. Within each area, we initially determined the Fano factor over a 100ms window, ensuring there was a 50ms overlap between successive windows. Following the mean-matching process, the survival rates of cells in these regions were observed to be notably high. Specifically, the aPFC region retained 84% of its cells, while the pPFC, MD, and VA regions had survival rates of 81%, 83%, and 82%, respectively. We found similar results with the mean-matched and raw (using all cells) Fano factor.

#### Pseudo-population neural decoding

To construct a robust and comprehensive training dataset and feature set for decoding purposes, we employed spike counts from each 5ms bin as input to our machine learning model. A pseudo-population of neurons was synthesized by: (i) incorporating neurons from all sessions into a consolidated vector, (ii) for each neuron/condition combination (with labels abstract rule, concrete rule, saccade direction), a random trial from the session where the cell was located and the aforementioned condition was present was selected without replacement, and (iii) this procedure was reiterated until we had chosen all trials for at least one session/neuron/condition. To effectively manage imbalances in the training data, the Synthetic Minority Over-sampling Technique (SMOTE) with a nearest neighbor approach was employed.

Our machine learning model is explicitly engineered to encapsulate both spatial (neuronal interactions) and temporal dependencies for the duration of a HRT trial, aligned to predetermined events (Abstract rule cue onset, Concrete rule cue onset, Saccade Onset). For extracting intricate spatial features from the spike counts of individual neurons and their combined patterns, we utilized a one-dimensional (1D) convolutional neural network (CNN) as the preliminary layer.

To ensure that the length of input to the succeeding layer remained constant, a max pooling layer was positioned after each CNN kernel. Our kernel dimensions were 3, 3, 5, and 7. Kernel sizes were meticulously chosen such that after passage through a 1×1 kernel, we had an outcome of 100 features as the input to our next machine learning module.

To account for the temporal dependencies of the firing pattern throughout the task, we employed a recurrent neural network (RNN) with a many-to-many architecture, which was designed to decode each condition using the enhanced features from the preceding layer. The proposed RNN comprises a Bidirectional Long-Short Term Memory (BiLSTM) layer, followed by a multiplicative local attention layer, a fully connected layer with dropout, and finally, a softmax layer for output. Given x(t) as the input at time t, the output of the BiLSTM forward path is computed as follows:

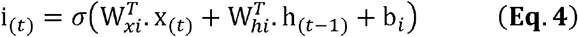

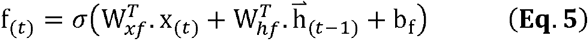

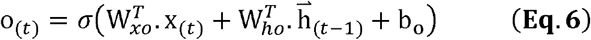

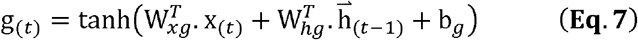

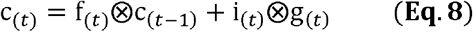

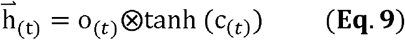

where i_(*t*)_,f_(*t*)_, c_(*t*)_, o_(*t*)_ are the input gate, forget gate, cell gate and output gate respectively. The weight matrices and bias vectors associated with the input, forget, cell, and output gates, as well as the hidden state at time *t*, are denoted by *W* (with appropriate subscripts) and *b* (with suitable subscripts). 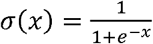 and 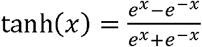, and the output of the backward path is 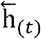

The attention layer output can be calculated as follows:

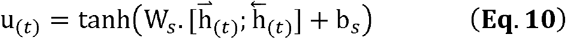

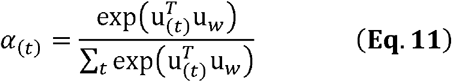

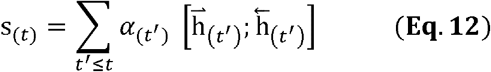

where:

- u_(*t*)_ is the activated attention weight at time *t*, computed through a tanh activation function applied to a linear combination of the forward and backward hidden states, 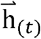 and 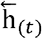 influenced by the attention weight matrix W_*s*_ and bias vector b_*s*_.
- *α*_(*t*)_ represents the normalized attention weight at time *t*, derived by applying a softmax function over u_(*t*)_ to ensure that the sum of the attention weights over all time steps equals one. This is achieved by exponentiating u_(*t*)_ scaled by u_*w*_ and then dividing by the sum of the exponentiated u_(*t*)_ values over all time steps, giving a probability distribution over the time steps.
- *s*_(*t*)_ stands for the context vector at time *t*, calculated as a weighted sum of the concatenated forward and backward hidden states up to time *t*, using the normalized attention weights *α*(_*t*′_).
- W_*s*_ is the weight matrix associated with the attention mechanism, which learns to weigh the importance of different parts of the hidden states for the computation of the attention weights.
- b_*s*_ is the bias vector associated with the attention mechanism, complementing W_*s*_ in learning the attention weights.
- u_*w*_ is a learnable parameter that scales u_(*t*)_ before the softmax operation in the computation of *α*_(*t*)_ learning to adjust the scaling of *u*_(*t*)_ to control the sharpness of the distribution.

The output of the attention layer was fed to a fully connected layer with dropout followed by a softmax layer which results in class conditional probability as equation 13 specifies.

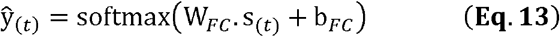

The weights and biases for all BiLSTM, attention, and fully connected (FC) layers undergo adjustment through backpropagation in time, a key algorithmic process integral to training our neural network model. The hyperparameters of the model, which include the quantity of hidden units in the BiLSTM, Attention, and FC layers, the learning rate, dropout probability, and the learning rate of stochastic gradient descent among others, are refined through an exhaustive grid search cross-validation.

In evaluating the efficacy of our model, decoding accuracy was ascertained using a 10-fold cross-validation method. Here, 60% of trials were used as the training set and the remaining 40% as the test set. The stratification and shuffling techniques were employed during this division of data sets, ensuring that we minimized any potential accuracy bias that could arise due to imbalanced classes.

To ensure a thorough and unbiased selection of trials, pseudo-population data generation was conducted 10,000 times. Accuracy measurements for test data were performed for each cross-validation fold and for each pseudorandom instance of the dataset, providing us with a robust statistical overview of the model’s performance.

The machine learning models were trained separately for each brain region of interest, including the aPFC, pPFC, MDc, and VAm. For the aPFC and pPFC in particular, we created four distinct models: one encompassing all cells and the remaining three models each representing superficial, middle, and deep layer cells.

#### Functional connectivity analyses

To measure the functional connectivity between neurons in PFC and thalamus, we applied the Adaptive Granger Causality (AGC) method ^39^. AGC is a dynamic and time-sensitive measure of directional interactions between neurons by modeling their spiking activities as a point process. Specifically, we used a conditional intensity function (CIF) to model the likelihood of neuronal spiking at any given moment, based on the spiking history of the neurons in the network. This approach captures the task-dependent dynamics of connectivity by linking the past activities of one neuron to the future activities of another. The AGC measure is derived using point process modeling, where the CIF describes the neuron’s firing probability at time t given its own spiking history and that of other neurons. This point process framework is well-suited for handling the non-Gaussian, binary nature of spike train data, making AGC a robust tool for capturing causal, directional interactions between neurons under varying task conditions.

We computed AGC over sliding 500 ms windows, stepped 100 ms, across a trial (Fig. 3F). In Fig. 3, G and H, we featured two windows following the onset of the abstract rule cue: 0-500 ms covering the cue presentation, and 500-1,000ms covering the delay period. The AGC was calculated for all bidirectional interactions between cell pairs for each trial. Once significant connections were identified using the Benjamini-Yekutieli procedure (described in the Quantification and Statistical Analysis section), we aggregated these interactions to the brain area level by summing the AGC values for all cell pairs between regions. For example, to compute the MDc-to-pPFC connectivity for a session, we summed the AGC values for all MDc-pPFC cell pairs and normalized the results for each time window. This allowed us to obtain an overall interaction strength between areas, averaged across all trials and normalized to ensure comparability across time windows. Finally, the 4×4 area-area connectivity matrices were averaged across all sessions for each 500 ms window, with the connectivity values normalized to range between 0 and 1.

#### Cortico-basal ganglia-thalamic model

To understand the underlying mechanism supporting the selection and maintenance of abstract rules, we built a layer-specific Cortico-Striatal-Thalamic model. Here, PFC includes three layers – superficial, middle, and deep. The trial starts with the abstract cue presentation (450 ms) followed by a delay (900 ms). Next, the concrete cue (450 ms) is presented, then a second delay (900 ms) and the three possible targets. As for the HRT, there were four cues that signal two abstract rules (shape or orientation) and nine concrete cues (from the combination of three shapes with three orientations). Each concrete cue shape and orientation mapped a response rule onto one of the three targets (contraversive, vertical, ipsiversive) depending on the currently relevant abstract rule. The blue triangle (BT) or red circle (RC) signals the shape rule, whereas the green circle (GC) or yellow triangle (YT) signals the orientation rule. The nine concrete cues consist of the three shapes (bowtie, rectangle and oval), each presented at three orientations (southwest, north, southeast). This model simulates the flow of information through the PFC layers, striatum, substantia nigra pars reticulata and thalamus. Specific parameter, network connectivity and synaptic weight values are shown in table S1, table S2 and table S3.

##### Leaky integrate-and fire-model

We used the leaky integrate-and-fire model (LIF) to simulate the activity of individual neurons. LIF is a simplified spiking neuron model where the electrical properties of a neuron’s membrane potential can be modeled with a parallel capacitor and a resistor. In this model, a neuron will fire when its membrane potential surpasses a certain threshold (−50 mv); after firing, its potential will reset (−65 mv) and cannot fire for a refractory time (1 ms). The time evolution of membrane potential (V) for neuron “j” is shown in Equation 14.

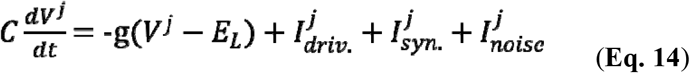

In this equation, we are computing how the voltage across the neuron “j” membrane changes as a function of its intrinsic driving current (I_driv._), synaptic inputs from its presynaptic cells (I_syn._) and stochastic inputs (I_noise_). Here, “*C”* is the membrane capacitance (1 nF), “t” is time, “E_L” represents resting potential (−65 mv), and “g” is the membrane conductance (g = 1, for simplicity). The neuron time constant is 20 ms.

The synaptic current (I_syn._) to the neuron “J” could be described with the general form represented in Equation 15, as follows:

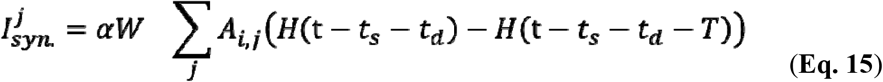

“W” stands for the synaptic weight of the incoming input. “A_i,j_ “is the adjacency matrix that determines from which neurons “*i*” there are incoming connections to the neuron “*j*.” Here, “H” is the *Heaviside* function controlling the synaptic current duration, “t_s_” is the time of presynaptic spike from the neuron “i,” “t_d_” is the synaptic delay between the two neurons (5 ms in most cases). “T” is the duration of the pulse that the presynaptic cell sends to the postsynaptic cell (1.5 ms for excitatory driving currents and 100 ms for modulatory effect). Here, “α” represents the gain modulation factor from VA to PFC superficial cortical cells (“α” = 2). This factor is 1 for other connections. When the pre-synaptic neuron is inhibitory, this synaptic current has a negative sign, and the negative pulse drops exponentially to compensate for the burst-like activity of fast-spiking cells (Equation 16).

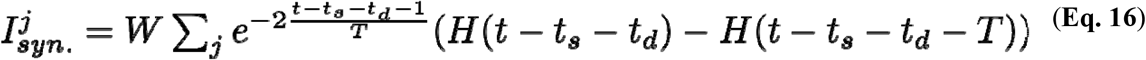

##### Network structure

###### PFC layers

For the abstract cue portion of the task, the first cells in the PFC that receive sensory signals and fire are the excitatory neurons in the middle layer (the middle layer consists of 50 neurons), which pass it to the superficial layer. Different populations of neurons in the middle layer get excited depending on the presented stimulus (e.g., BT vs. GC). For simplicity, we modeled starting cells in the superficial layer, which receives the sensory signal from the middle layer, in four chains (each chain has 50 neurons, a total of 200 superficial PFC cells) associated with each visual stimulus (BT, RC, GC and YT). Each neuron is connected locally to the neurons within and across the chains; connections within the chains are 1.3* stronger than across the chains. These superficial cells are also connected to a population of secondary PFC cells that get activated (and become rule-selective) later (the superficial layer consists of 100 of these rule-selective ensemble neurons). These four superficial chains converge to four chains in the deep layer (similar to the superficial layer, each chain has 50 neurons, for 200 total deep layer PFC neurons) with a connectivity matrix that has the strongest weight on the diagonal (*\*2*, from each superficial chain to its mirror chain in the deep layer, for example, from *superficial chain 1* [BT] to *deep chain 1*) and weaker weights off the diagonal (*\*0*.*2* from superficial chain 1 [BT] to deep chain 2, 3, 4). To investigate the effects of convergence from superficial to deep cortical layers, we also adjusted the weights off the diagonal that corresponded to the same rule (*\*0*.*2 and greater* from superficial chain 1 [BT] to deep chain 2 [RC]; *0.7 and *0.8 in fig. S12). This suggested that the range of plausible convergence from superficial to deep cortical layers does not generate rule-selectivity in PFC early after abstract cue onset. Although stronger convergence might eventually lead to early, abstract rule-selectivity in the cortex, it would not be consistent with the electrophysiological data (e.g., Fig. 2L).

A similar structure is assumed for the concrete cue portion of the task, except the PFC superficial layer has six chains (each chain has 50 neurons, a total of 300 superficial PFC cells) that will respectively respond to one of three shapes or one of three orientations. Additionally, each PFC concrete rule cell type is associated with a specific choice (there are 50 neurons representing each choice, for a total of 150 choice neurons). As a result of abstract cue processing, when the PFC abstract rule ensemble cells fire, it stimulates inhibitory cells (there are 100 inhibitory neurons) that inhibit the PFC superficial concrete rule cells that represent the irrelevant concrete rule class. For example, firing of the PFC shape rule ensemble will inhibit the three groups of PFC superficial orientation concrete rule cells (which respond to the north, southwest and southeast orientations). Subsequently, the concrete cue appears (e.g., northward-oriented bowtie), and each concrete cue has two features: shape and orientation. So the cue (via activation of excitatory neurons in the middle layer (50 neurons)) can stimulate two PFC superficial concrete rule cell types that match the cue shape and cue orientation, respectively. However, since there is always one abstract rule/feature that is currently relevant, the activated superficial PFC abstract rule ensemble (e.g., shape) inhibits the concrete rule cells (e.g., SW, N, SE) which correspond to the irrelevant abstract rule (e.g., orientation). Because of this, only one of the PFC superficial concrete cell types (e.g., bowtie) will stimulate the corresponding MD action choice cells (e.g., contraversive).

###### Disinhibition of VA

The PFC deep layer projects to the striatum, modeled with a population of inhibitory neurons. Upon the presentation of the abstract cue, an elevated activity of the PFC deep layer causes firing in the striatum (the striatum consists of 50 neurons), inhibiting the inhibitory cells in the SNr (the SNr consists of 50 neurons). As SNr targets VA, this temporary decrease in SNr activity creates an excitable window for VA activity.

###### VA and MD thalamus

We modeled VA and MD thalamic nuclei. VA and MD cells integrate the synaptic inputs from their pre-synaptic deep layer PFC cells. For the abstract rule portion of the task, the first two chains (1 and 2) from deep PFC converge onto VA and MD shape-selective cells (VA and MD consists of 50 shape-selective neurons), and the second two chains (3 and 4) project onto MD and VA orientation-selective cells (VA and MD consists of 50 orientation-selective neurons). To be consistent with the known anatomical features of these circuits, in this model, connections projecting to the VA have stronger weights (*5) than those targeting MD. As mentioned above, the PFC-striatum-driven disinhibition of SNr allows for a facilitating time window for VA firing. The synchronous input from deep PFC recruits the VA and MD thalamic cells. VA gets activated first (t_d_ = 5 ms), as the time needed to reach MD is longer (t_d_ = 10 ms), due to the extra anatomical connection from cortical layer 5 to 6 (i.e., matrix cells, e.g., in VA, receive input from layer 5; and core cells, e.g., in MD, receive input from layer 6) and weaker modulatory PFC inputs. VA activation sends a broad pulse to their post-synaptic PFC cells, including the secondary PFC cells. VA cells have divergent modulatory (amplifying the gain of cortico-cortical connections by factor two) and excitatory effects on their post-synaptic PFC cells. The “secondary” population of PFC cells will receive driving and modulatory input from VA and driving current from MD. After activation, this population of PFC cells recruits more MD cells, which creates a PFC-MD loop to support rule information maintenance during the delay period. We added extended delay time within this PFC-MD loop to consider extra connections across layers that we are not modeling. The concrete cue portion of the task is modeled similarly to the abstract cue portion for MD, but MD is modeled with three cell groups corresponding to the contraversive, vertical and ipsiversive choice options (each group consists of 50 neurons, a total of 150 MD cells). The activated MD cells will stimulate the corresponding PFC choice cells, which further stimulate MD. They will form a reverberating effect similar to the abstract cue cases. However, the PFC ensemble representing the choice does not need stimulation from VA to recruit enough cells to support the maintained activity between MD and PFC choice cells.

###### Cortical inhibition

For the abstract rule, in addition to VA’s excitatory impact on superficial PFC cells, it modulates the connections within the PFC superficial layer and drives PFC inhibitory cells (there are 50 such PV neurons), resulting in a general inhibition. A more specific inhibition happens between the competing PFC *shape* and *orientation* (“secondary”) neuronal ensembles. MD drives cortical inhibitory cells (there are 100 such FS neurons), which target the irrelevant rule modality. This inhibition creates the “*winner-take-all*” paradigm that protects the fidelity of the signal. For the concrete rule, MD drives the “winner-takes-all” inhibition (there are 150 FS inhibitory neurons) to inhibit the other two irrelevant PFC choice ensembles (e.g., MD left choice cells will inhibit the PFC vertical choice ensemble and PFC ipsiversive choice ensemble).

###### Lesions of VAmc or MDpc

We model lesions of VA and MD by disconnecting them from the rest of the network.

#### Decoding Simulated Data

To decode abstract rules from simulated data in each of three conditions — namely, “healthy”, MD lesion, and VA lesion — we employed the same machine learning model previously used for decoding of electrophysiological data. This model operates with a 50 ms time window for counting spikes. However, the method for generating the pseudopopulation of cells was adjusted.

For the simulated data, we randomly selected between 50% and 75% of cells at each step of randomization. Specifically, we chose a random integer from 50 to 75 to represent the percentage of cells to be selected. This selection was uniformly made from each area and under each abstract rule to ensure data balance. Decoding was then carried out using 10-fold cross-validation for each random iteration. This process was repeated for 10,000 iterations across time, resulting in the generation of accuracy metrics.

### QUANTIFICATION AND STATISTICAL ANALYSIS

#### Wilcoxon rank-sum tests for selectivity analyses of spike data

We aimed to ascertain the proportion of selective cells in various regions including aPFC, pPFC, MD, and VA. To this end, we first calculated the spike counts for each defined interval of interest. Subsequently, we carried out a non-parametric Wilcoxon rank-sum test for every cell in each specific condition, namely, Abstract Rule, Concrete Rule, Saccade Direction, and Outcome.

To mitigate the challenges arising from multiple comparisons, we applied Holm’s correction. A cell was classified as “selective” based on a final p-value less than 0.05, following the adjustment for multiple comparisons.

Furthermore, we determined the proportion of neurons exhibiting selectivity for the abstract rule condition over time, employing 100 ms running windows with a 50 ms overlap. This analysis invoked the same statistical test as previously described. To substantiate the significance of the calculated proportions, we executed a 10,000-step bootstrapping process, wherein we randomly chose 80% of the cells and assessed the proportion of selective cells in each iteration. Moreover, we contrasted the selectivity against a null distribution, derived from permutating the abstract rule labels and determining the selective cell proportions in each permutation.

To calculate the latency, we measured the times when the portion of selective cells reached 50% of their maximum selectivity across all 10,000 bootstrap samples. Then, we conducted a linear trend analysis using a General Linear Model (GLM) and linear contrasts

##### Statistical analyses of population SI timecourse

We compared the timing and magnitude of cognitive information represented by single neurons in the PFC and thalamus. To do this, we ran a univariate sliding window analysis (100 ms window width, stepped by 100 ms) of the population SI curves. Based on the thalamo-cortical model in Fig. 1G that we envisioned supporting accurate performance of the HRT, we hypothesized that thalamic neuron populations would have stronger abstract rule-selectivity than PFC populations (due to convergent inputs to the thalamus amplifying weak input signals). We used one-tailed t-tests to probe whether thalamic mean abstract rule SI values were greater than PFC mean SI values in 100 ms-wide nonoverlapping windows of the SI curves beginning 100 ms prior to abstract rule cue onset and continuing until the window beginning 800 ms after cue onset (corresponding to the shortest duration of delay 1, and common to all trials). For the concrete rule period, we used two-tailed t-tests, because we hypothesized that intracortical mechanisms and convergent cortico-thalamic mechanisms could give rise to different magnitudes of concrete rule SI. We applied the same sliding window analysis for this period, as well. Finally, for the saccade aligned analysis of directional-selectivity, we applied the same analysis to the three time windows: pre-saccade direction SI windows spanned from 700 ms until 100 ms prior to saccade onset; peri-saccade direction SI windows spanned from 100 ms prior to saccade until 100 ms following saccade onset; and post-saccade direction SI windows spanned from 100 ms until 900 ms post saccade onset.

#### Statistical analyses of decoding data

We scrutinized the decoding results of each ROI at each time bin. This was accomplished by comparing these results to a null distribution. This null distribution was derived through the permutation of class labels 10,000 times. Following this, we computed the accuracy at each instance to establish the null distribution.

To draw comparisons between the accuracies of different brain regions, a two-step statistical process was employed. Initially, the existence of an effect was ascertained using an analysis of variance (ANOVA). The ANOVA results, specifically F-statistics and p-values, were reported as effect size when necessary.

Post this, for each hypothesis a detailed contrast analysis was designed and conducted for each time bin. Below is an example of contrast design for one of the hypotheses: To rigorously test the specified hypotheses regarding the conveyance of abstract rule information among the various ROIs, namely VAm, MDc, pPFC and aPFC, the following series of contrasts was designed and implemented.

For the first hypothesis positing that the VAm population carries more abstract rule information than other areas, the contrast was structured as follows: [VAm - (MDc + pPFC + aPFC)/3]. This contrast is representative of the differential encoding of abstract rule information between VAm and the mean of MDc, pPFC and aPFC. A positive value from this contrast would support the hypothesis, suggesting that the VAm carries more abstract rule information than the other regions.

The second hypothesis asserts that the MDc population carries more abstract rule information than both the pPFC and aPFC. This was investigated through the following contrast: [MDc - (pPFC + aPFC)/2]. This contrast represents the difference in abstract rule information encoding between MDc and the mean of the pPFC and aPFC. A positive value here would support the hypothesis, suggesting that the MDc carries more abstract rule information than the pPFC and aPFC.

Lastly, for the third hypothesis stating that the pPFC carries more abstract rule information than the aPFC, we utilized the contrast [pPFC - aPFC]. A positive value from this contrast would support the hypothesis, indicating that the pPFC encodes more abstract rule information than the aPFC.

In more succinct notation, the designed contrasts for hypotheses tested and reported in the main text for abstract rule decoding are:

For the hypothesis that the VAm population carries abstract rule information earliest, the contrast is set as: [0, 0, 0, 1] for [aPFC, pPFC, MDc, VAm] respectively.

1. For the hypothesis that thalamic populations VAm and MDc carry rule information earlier than pPFC, the contrast is set as: [0, -1, 1, 1] for [aPFC, pPFC, MDc, VAm] respectively.
2. For the hypothesis that the VAm population carries more abstract rule information than the other ROIs 0-450 ms after abstract rule cue onset, the contrast was set as: [-1, -1, - 1, 3] for [aPFC, pPFC, MDc, VAm] respectively.
3. For the hypothesis that MDc carries more abstract rule information than both pPFC and aPFC 0-450 ms after abstract rule cue onset, the contrast was designed as: [-1, -1, 2, 0] for [aPFC, pPFC, MDc, VAm] respectively.
4. For the hypothesis that pPFC carries more abstract rule information than aPFC 0-450 ms after abstract rule cue onset, the contrast was set as: [-1, 1, 0, 0] for [aPFC, pPFC, MDc, VAm] respectively.
5. Each of these contrasts, carried out for each time bin, were corrected for multiple comparisons using the Bonferroni method. These results were then reported with the respective t-statistics and p-values where necessary, highlighting the evidence supporting these hypotheses.

Finally, the latency for each ROI was calculated. This was accomplished by utilizing the decoding accuracy results and determining the time at which each region possessed decodable information, i.e., significantly above chance levels, as compared to the null accuracy distribution.

#### Statistical analyses of functional connectivity data

Given the large number of potential connections when calculating AGC for all bidirectional interactions between cell pairs, we applied a False Discovery Rate (FDR) correction to limit the rate of false positives. Specifically, we used the Benjamini-Yekutieli (BY) procedure, which is an enhancement of the standard FDR correction. Unlike traditional methods, the BY procedure accounts for dependencies between tests, providing more stringent control over false discoveries, especially in high-dimensional data such as neuronal connectivity networks. The BY test allows us to identify statistically significant interactions by controlling the expected proportion of false positives among the rejected hypotheses. This is particularly crucial in our analysis, where we are testing for multiple connections across different time windows. By employing the BY correction, we ensure that the set of significant connectivity links is reliable.

We reported the directional connectivity values between the key regions during abstract rule processing, i.e., MDc-to-pPFC, pPFC-to-MDc, VAm-to-pPFC and pPFC-to-VAm. To statistically compare connectivity measures in each direction, we used t-tests to assess the differences in connectivity between area pairs.

