## Supplementary materials for "Primate thalamic nuclei select abstract rules and shape prefrontal dynamics"

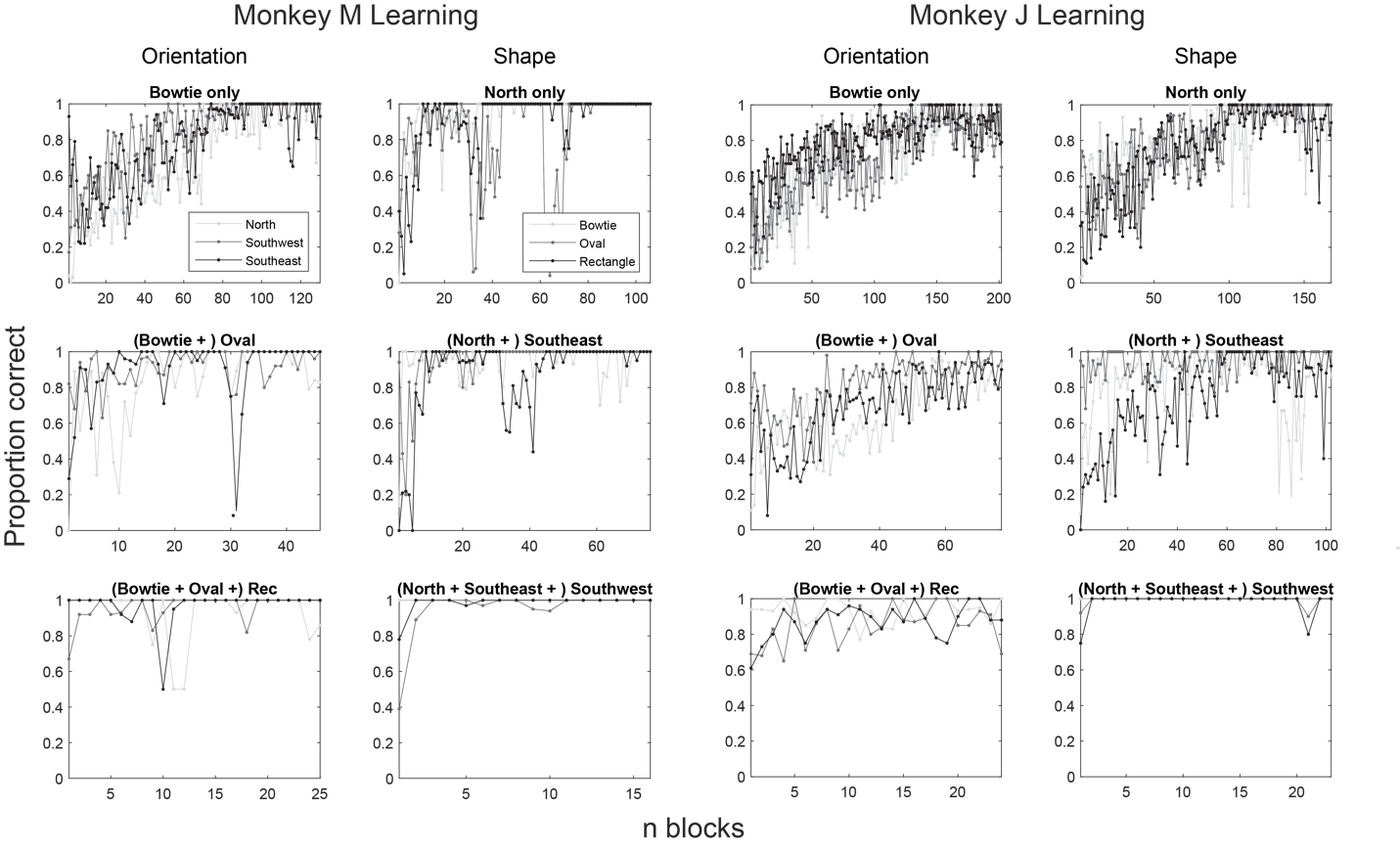

**Figure S1. Learning curves for the two monkeys during training on the HRT. Related to Figure 1.** Under each abstract rule, monkeys began training with concrete cues associated with only one irrelevant feature that varied only in the relevant dimension (e.g., bowtie as the single irrelevant shape feature while training on the orientation rule set with all three orientations). As is typical for non-human primate learning of cognitive tasks, learning occurred slowly through operant mechanisms (see top row). However, when concrete cues associated with a second irrelevant feature were added, learning occurred much more quickly (see second row). When the cues associated with the final irrelevant feature were added, generalization of the rule set occurred immediately (see third row). Datapoints represent accuracy measured over the course of 100-trial blocks.

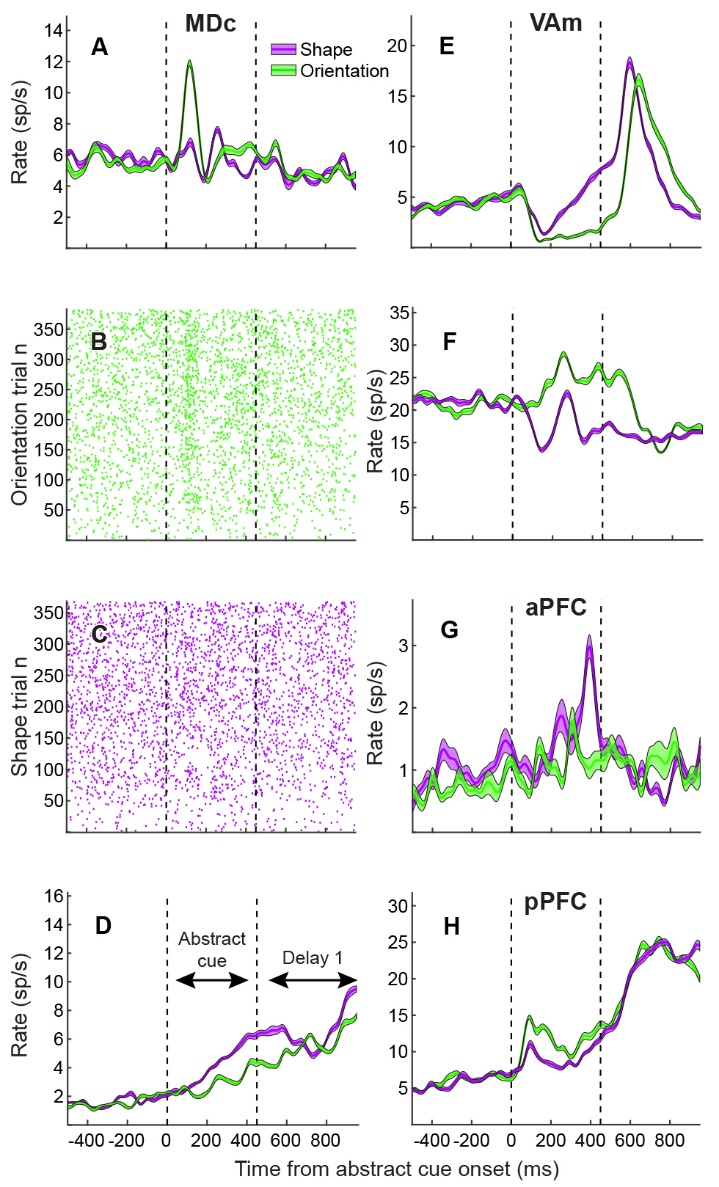

**Figure S2.** **Comparison of abstract-rule selective neurons in thalamus and cortex. Related to Figure 2.** MDc SDF (**A**) and corresponding raster plots for orientation (**B**) and shape (**C**) showing abstract rule selectivity at onset. (**D**) MDc neuron showing sustained selectivity. (**E** and **F**) Two VAm neurons showing selectivity starts early. Selective cortical neurons in aPFC (**G**) and pPFC (**H**). Although the aPFC neuron in (G) shows rule-selectivity, abstract-rule selective neurons were less common in aPFC compared with other areas (Fig. 2J).

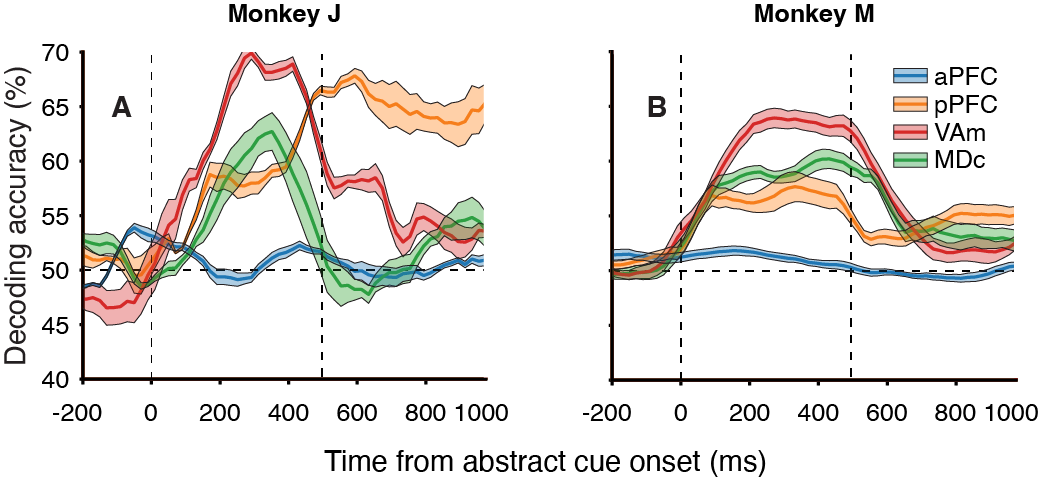

**Figure S3. Single subject abstract rule decoding. Related to Figure 2.** Timecourses of rule (shape vs orientation) decoding accuracy for each ROI, aligned to abstract rule cue onset (50% accuracy represents chance performance), for monkey J (**A**) and monkey M (**B**). Timecourses of abstract rule decoding accuracy and other data for each monkey were similar, which motivated combining their data for the remainder of analyses.

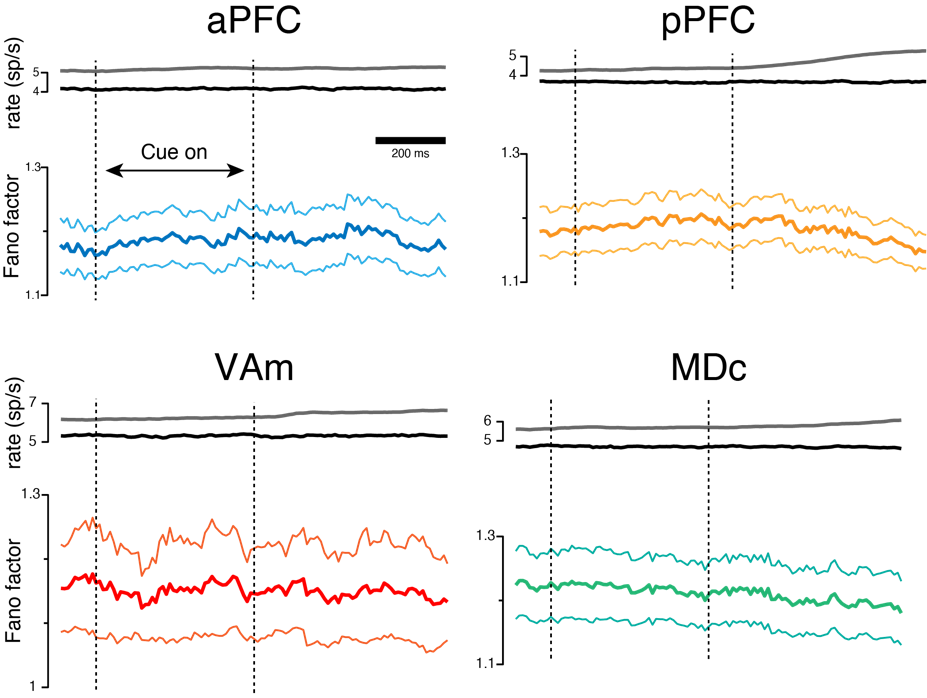

**Figure S4. Fano Factor does not differ between ROIs. Related to Figure 2.** Population mean firing rate after the mean-matching process in black, at top of each ROI’s panel; raw mean firing rate, prior to mean-matching, in grey. Mean-matched Fano factor calculated in sliding 100-ms windows, with 50-ms overlap between successive windows, at bottom of each ROI’s panel. Firing rate and Fano factor from 100 ms before to 1000 ms after abstract rule cue onset.

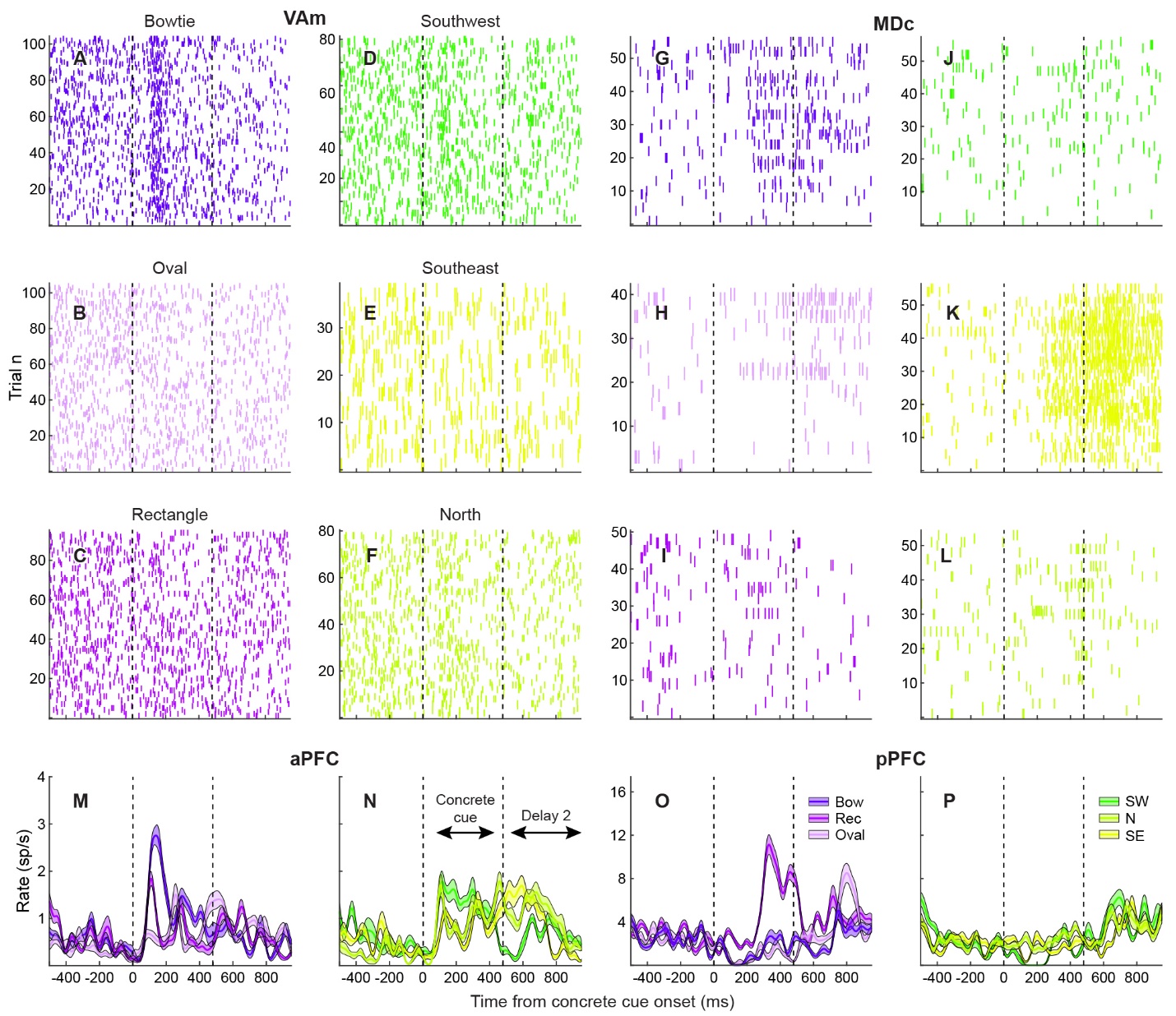

**Figure S5. Comparison of concrete-rule selective neurons in thalamus and cortex. Related to Figure 4.** Raster plots associated with single neuron examples in Fig. 4, A and B (**A** to **F**) and Fig. 4, C and D (**G** to **L**). SDFs of neurons in aPFC (**M** and **N**) and pPFC (**O** and **P**).

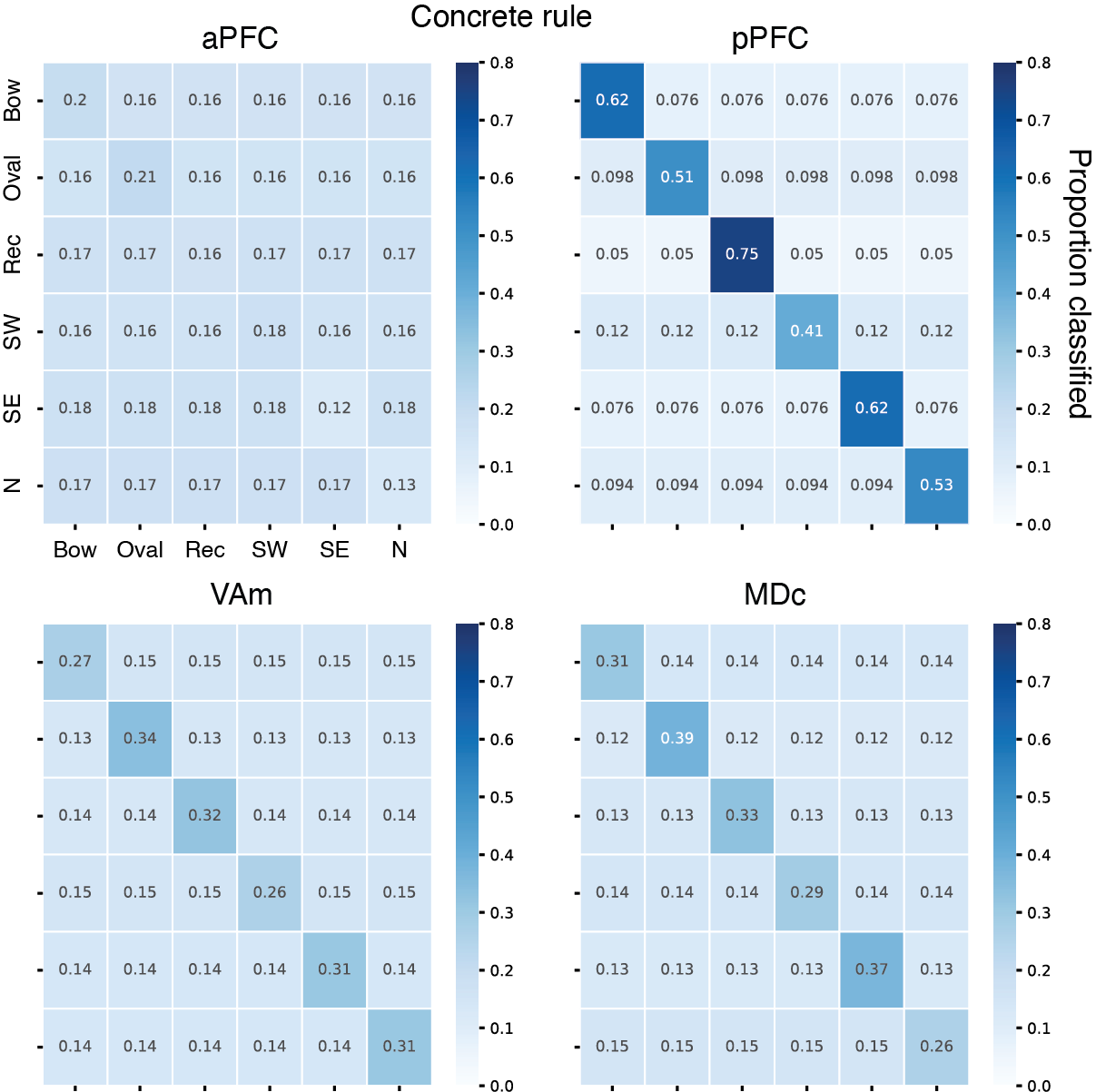

**Figure S6. Confusion matrices for concrete rule decoding. Related to Figure 4.** Decoding accuracy color-coded. Darker diagonal indicates decoding of each of the six concrete rules.

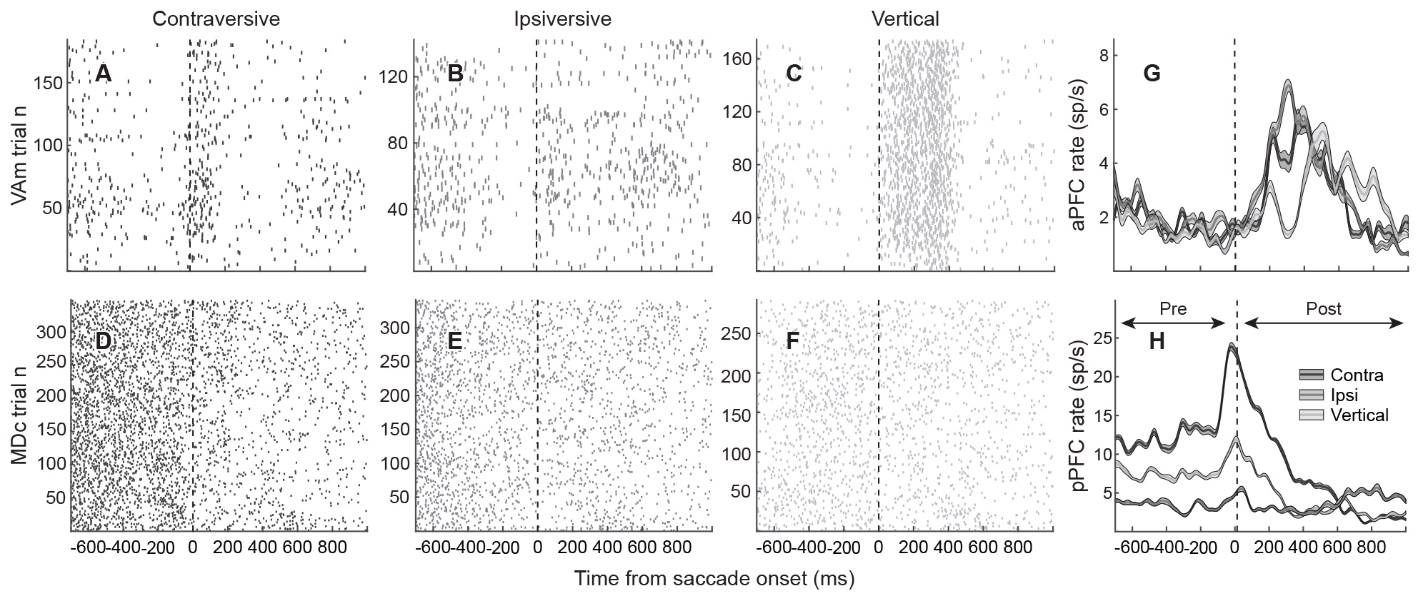

**Figure S7. Comparison of direction-selective neurons in thalamus and cortex. Related to Figure 5.** Raster plots associated with single neuron examples in Fig. 5A (**A** to **C**) and Fig. 5B (**D** to **F**). SDFs of aPFC (**G**) and pPFC (**H**) neurons.

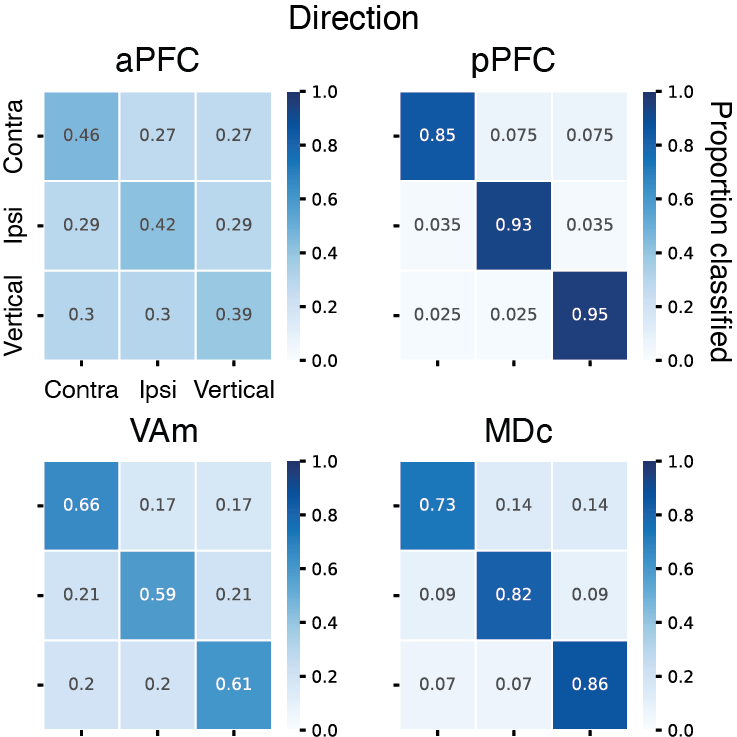

**Figure S8. Confusion matrices for direction decoding. Related to Figure 5.** ­Darker diagonal indicates decoding of each of the three directions.

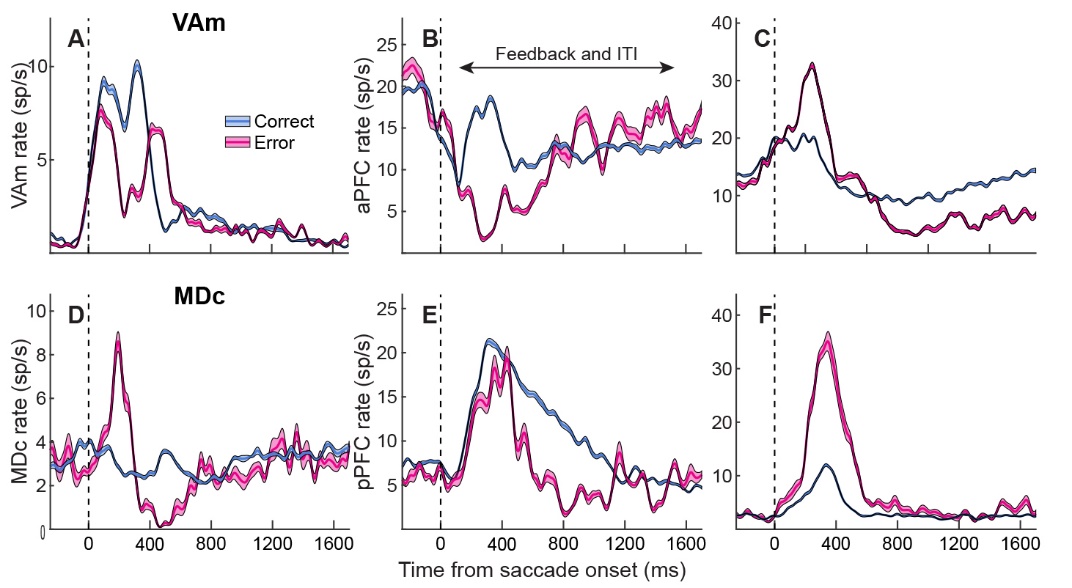

**Figure S9. Selectivity for trial outcome in thalamus. Related to Figure 6.** SDFs of VAm (**A** to **C**) and MDc (**D** to **F**) neurons.

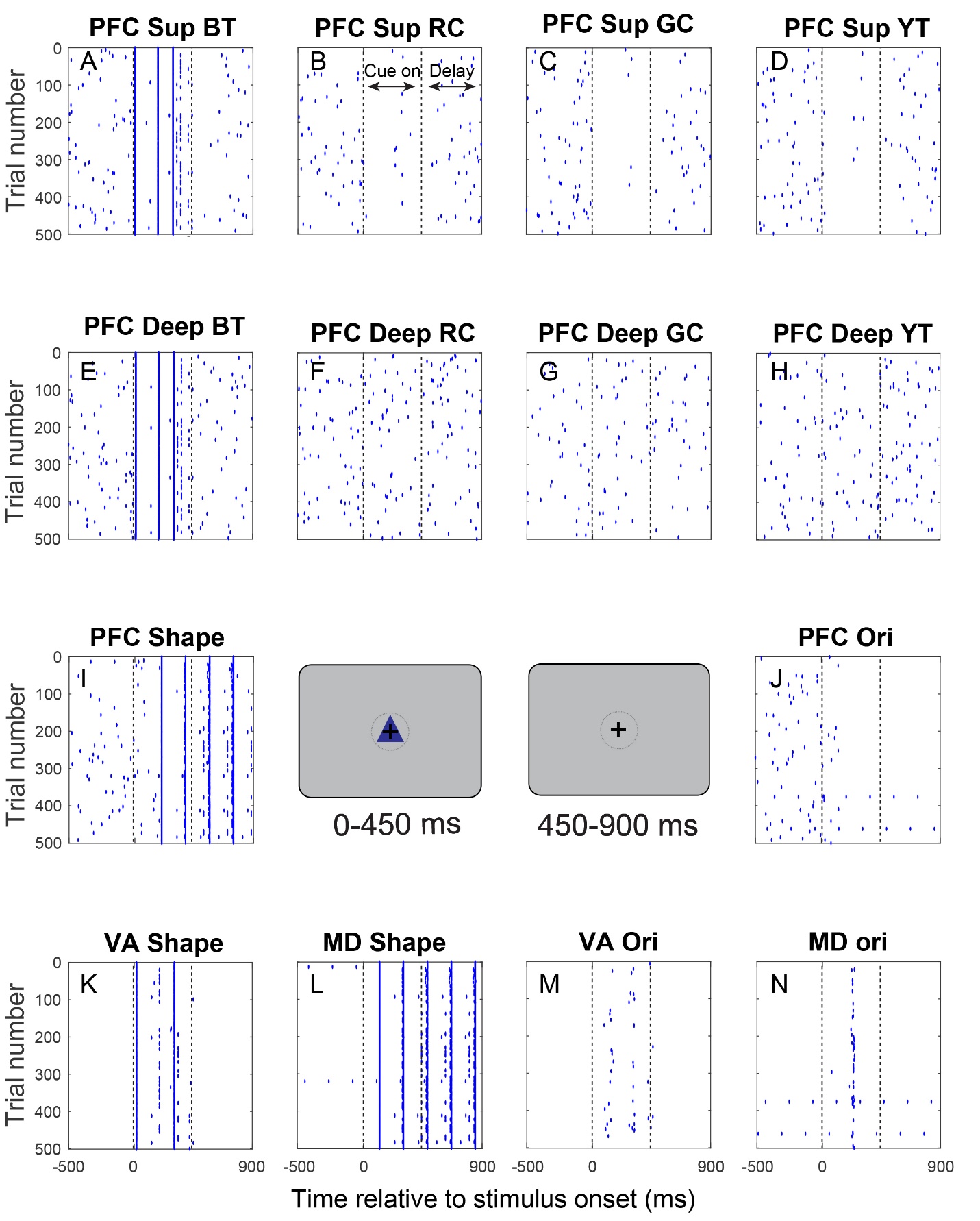

**Figure S10. Model spike output (rasters) for blue triangle cue (shape rule) trials. Related to Figure 7.** Each raster corresponds to individual cell from model populations described in Fig. 7A. Insets represent modeled HRT trial epochs: visual stimulus (abstract rule cue) and delay period. Sup, superficial; BT, blue triangle; RC, red circle; YT, yellow triangle; GC, green circle; Ori, orientation.

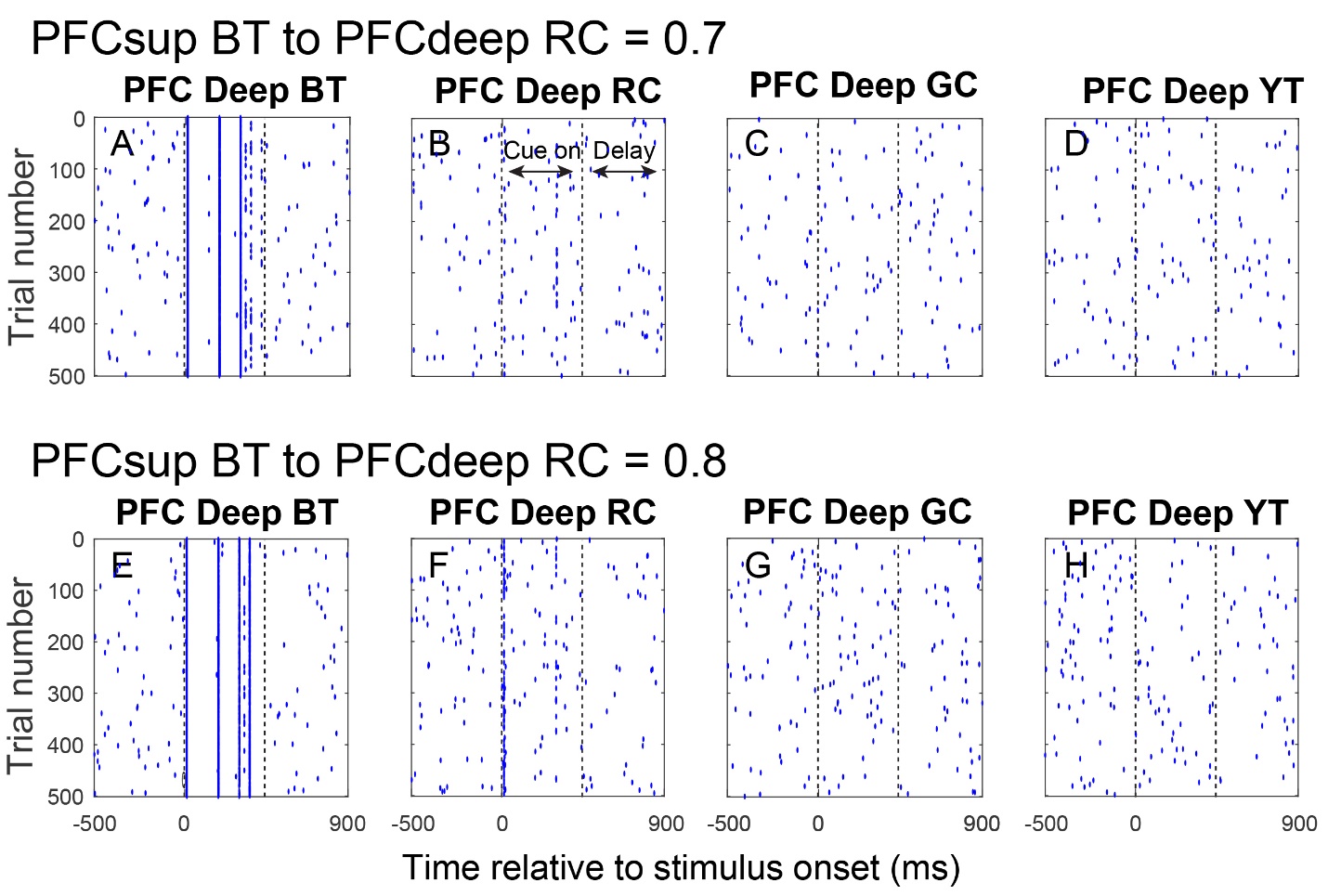

**Figure S11. Increasing convergence from PFC superficial layers to PFC deep layers *in silico* does not generate early abstract rule selectivity in PFC, in line with *in vivo* results. Related to Figure 7.** Model spike output (rasters) for PFC deep layer cells shown for two different model configurations, on blue triangle cue (shape rule) trials. Each row corresponds to a model with a different synaptic weight (convergence) from PFC superficial BT cells to PFC deep RC cells. For comparison, the model configuration in fig. S9 has four populations of PFC superficial cells (BT, RC, GC, YT) converging on to four populations in the deep layer, with a connectivity matrix that has the strongest weight on the diagonal (**2* from the superficial BT population to the deep BT population) and weaker weights off the diagonal (**0.2* from the superficial BT population to deep populations RC, GC and YT). Here, in fig. S10, the model configurations have adjusted weights off the diagonal that correspond to the same rule: (top row) **0.7* from the superficial BT population to the deep RC population; and (bottom row) **0.8* from the superficial BT population to the deep RC population. BT, blue triangle; RC, red circle; GC, green circle; YT, yellow triangle.

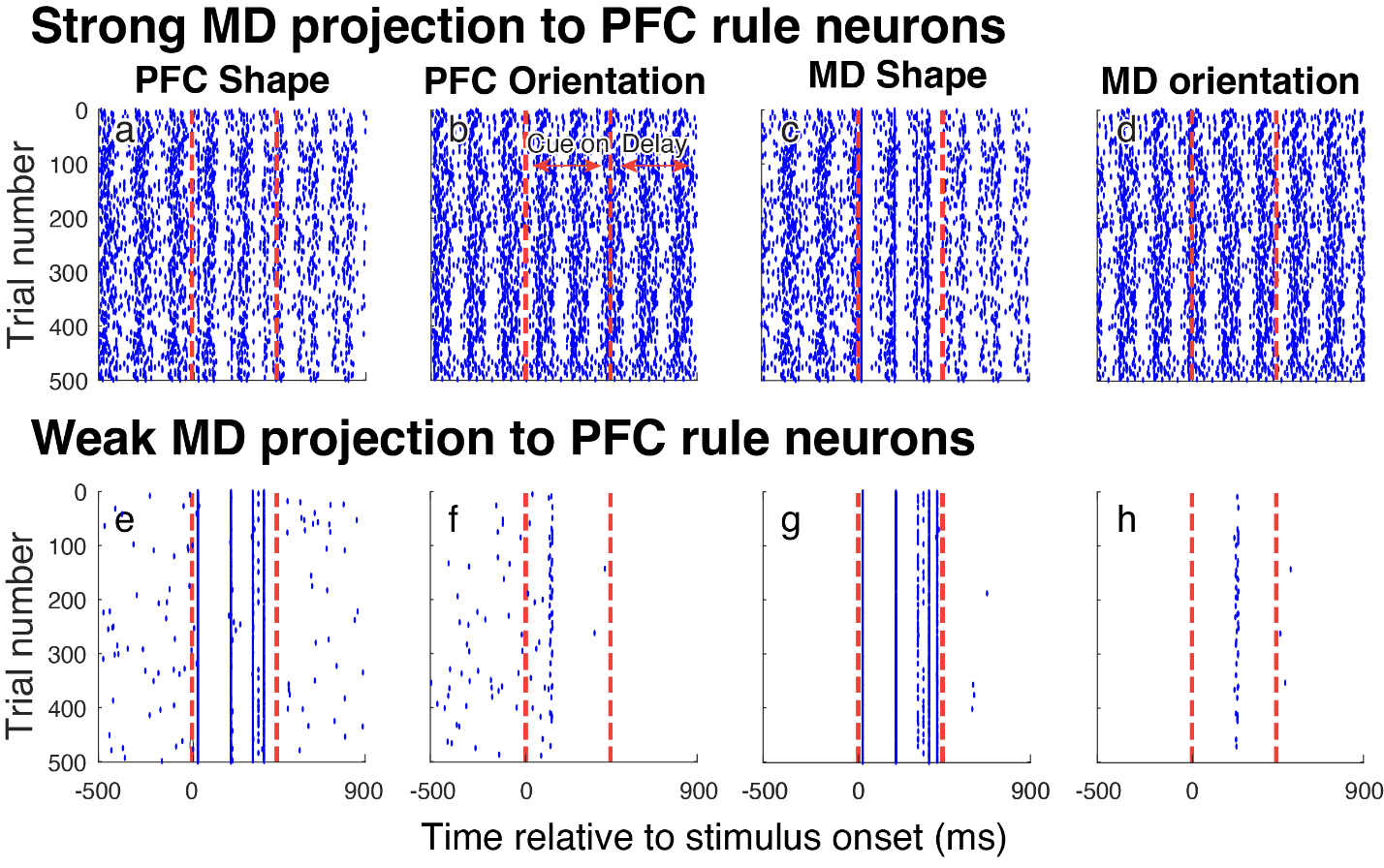

**Figure S12. Compared to “healthy” model, stronger weight of MDpc projection to PFC rule-selective neurons increased activity for both abstract rules (thereby reducing rule-selectivity); and weaker weight of MDpc projection to PFC rule-selective neurons decreased delay period rule-related activity. Related to Figure 7.** Model spike output (rasters) for PFC rule cell shown for two different model regimes, on blue triangle cue (shape rule) trials. Each row corresponds to a model with a different synaptic weight from MDpc rule cells to PFC rule cells: stronger weight in top row, and weaker weight in bottom row, compared with the original weighting in the “healthy” model (our baseline regime). For the strong connectivity regime, the new weighting is 0.036 (10x the original synaptic weight); for the weak connectivity regime, the new weighting is 0.00072 (0.2x the original synaptic weight).

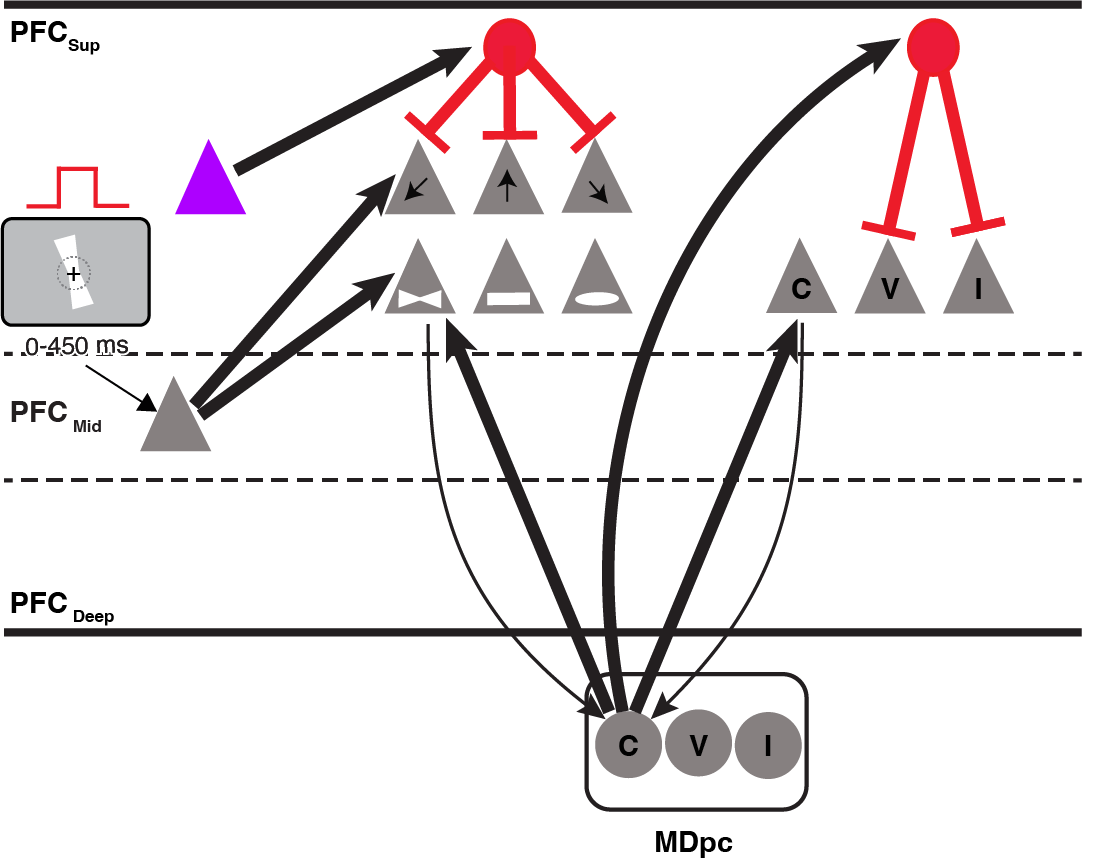

**Figure S13. Extended model for concrete rule and choice representation. Related to Figure 7.** The model in Fig. 7 processes the abstract rule, and the abstract-rule selective cell populations in the PFC superficial layer in Fig. 7 (corresponding to the purple triangle here in fig. S13) provide input to a population of inhibitory neurons in the extended model here in fig. S13. The extended model processes the concrete rule and outputs a choice. The *in silico* PFC includes superficial and middle (concrete rule cue input layer) layers; but cells in the deep layer in this extended model component are not explicitly modeled, instead they are accounted for in the transmission time from PFC to MDpc. MDpc receives modulatory input from PFC; and MDpc provides driving input to PFC, reflecting known primate anatomy and physiology ^8^. Flow of information through PFC and MDpc simulated using leaky integrate-and-fire cells. Each triangle or circle represents a different cell population. PFC superficial layer contains populations selective for one of the six concrete rules (SW, N, SE, bowtie, rectangle, oval) or one of the abstract rules (purple, shape rule; green, orientation rule (not shown)). These abstract rule-selective cell populations correspond to the abstract rule-selective cell populations in the PFC superficial layer in the model shown in Fig. 7. A population of inhibitory interneurons, driven by the relevant abstract rule-selective cell population (in this case, shape, as cued by a blue triangle stimulus) in the PFC superficial layer, imposed inhibition on the subset of concrete rule-selective cell populations (in this case, SW, N and SE) in the PFC superficial layer, which correspond to the irrelevant abstract rule (in this case, orientation). A second population of inhibitory interneurons, driven by MDpc, specifically inhibited superficial cells selective for the irrelevant saccade targets (in this case, vertical and ipsi). MDpc cell populations shown are selective for the contra (C), vertical (V) or ipsi (I) saccade targets, reflecting known subpopulations of motor-related MD neurons in primates ^17^. Black arrows represent excitatory projections, and red lines with flat terminations indicate inhibitory projections (model connectivity in table S1). Thicker black lines represent excitatory projections having synaptic weight > 0.003; thinner lines represent excitatory projections having synaptic weight ≤ 0.003 (model synaptic weights in table S3).

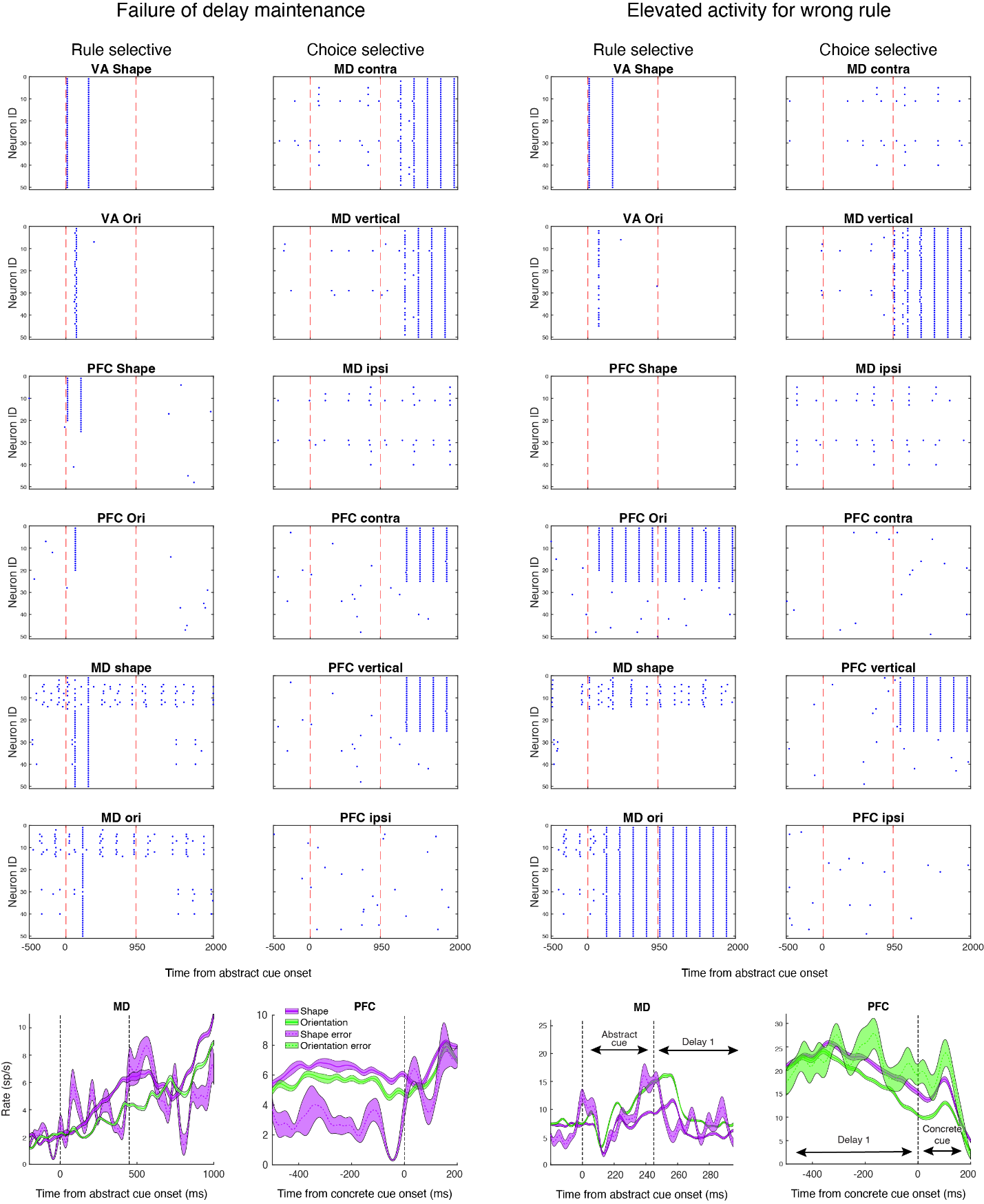

**Figure S14. *In silico* cell activity on error trials resembles *in vivo* single cell activity on error trials. Related to Figure 7.** Model spike output (rasters) for trials where a blue triangle cue (indicating the abstract shape rule is relevant) was presented followed by a northward-oriented bowtie cue (indicating the concrete bowtie rule is relevant, and the contra target is the correct choice). Far-left and left columns show rasters for an error trial where there was little delay activity in PFC and MD after the abstract rule cue presentation. Right and far-right columns show rasters for an error trial where there was more activity for the wrong rule. Each raster corresponds to 50 cells from model populations described in Fig. 7A (abstract rule processing) and fig. S13 (concrete rule and choice processing). First and second red dashed vertical lines respectively show onset of abstract rule cue and concrete rule cue. Bottom row shows *in vivo* example of reduced delay activity in error trial for MD cell (far left) and PFC cell (left), as well as example of more activity for the wrong/irrelevant abstract rule in error trial for MD cell (right) and PFC cell (far right).

| **Description** | **Value** |
| --- | --- |
| Neuron time constant | 20 ms |
| Step time size | 0.01 ms |
| Synaptic delay | 5 ms |
| MD synaptic delay | 10 ms |
| Firing threshold | -50 mv |
| Neuron resting potential | -65 mv |
| Membrane resistance | 10 Ω.cm^2^ |
| Membrane conductance | 1 nF |
| Refractory time | 1 ms |
| Pulse width for most PFC cells | 1.5 ms |
| Pulse width for general inhibition | 100 ms |
| Pulse width for winner-take-all | 1 s |
| Window width for VA | 100 ms |
|  | **Connectivity Patterns** |
| Middle layer PFC cells to superficial (abstract and concrete) | All-to-all connectivity in sequential blocks of ten; the last ten neurons are connected to the first ten neurons. |
| Within superficial PFC cells (abstract and concrete) | All-to-all connectivity in sequential blocks of ten, within and across chains; the last ten neurons are connected to the first ten neurons. |
| Within deep layer PFC cells (abstract only) | All-to-all connectivity in sequential blocks of ten, within and across chains; the last ten neurons are connected to the first ten. |
| Superficial to deep layer (abstract only) | Corresponding sequential blocks of ten converge onto the deep layer |
| Deep PFC to MD (abstract only) | All deep layer chains 1&2 converge onto the first block of MD shape (30%), and from 3&4 converge onto the first block of MD orientation (30%) |
| Deep PFC to VA (abstract only) | From PFC deep first two blocks (40%, chains 1 & 2/ chains 3 & 4) converge to all VA cells (shape/ orientation) |
| VA to PFC rule ensemble (driving) (abstract only) | VA to PFC 40% rule selective ensemble |
| VA to PFC superficial (modulatory) (abstract only) | 20% of VA cells projected back to all superficial PFC cells |
| MD to PFC (abstract and concrete) | All MD to 50% of PFC rule selective ensemble and MD randomly projects to the 10% superficial PFC cells |
| PFC rule ensemble to MD (abstract and concrete) | All to the same rule MD |
| PFC abstract rule ensemble to concrete inhibitory cells | All-to-all connectivity |

**Table S1. Model parameters and connectivity patterns. Related to Figures 7 and S13.**

|  | **Synaptic Weights** |
| --- | --- |
| PFC middle to PFC superficial | 0.02 |
| PFC superficial within chains (except rule ensemble) | 0.0012 |
| PFC superficial and deep across chains | 0.0009 |
| PFC superficial to deep within chains | 0.008 for the first 10 superficial neurons to first 10 deep neurons and 0.004 for the other neurons |
| PFC superficial to deep across chains | 0.0016 for the first 10 superficial neurons to first 10 deep neurons and 0.0008 for the other neurons. As shown in fig. S11, we also increased weights from PFC superficial to deep across chains representing the same abstract rule. See “PFC layers” subsection in “Network structure” section of Methods for details. |
| PFC deep to MD | 0.003 |
| PFC deep to VA | 0.015 |
| PFC rule to MD | 0.0018 |
| PFC rule within chains | 0.002 |
| PFC deep to ST | 0.0145 |
| VA to PFC rule | 0.01 |
| VA to PFC inhibitory cells | 0.0145 |
| MD to PFC rule | 0.0036 |
| MD to PFC superficial | 0.012 |
| MD to PFC inhibitory cells | 0.001 |
| Inhibitory cells to PFC rule | 0.0025 |
| ST to SNpr and SNpr to VA | 0.001 |
| Inhibitory cells to PFC superficial | 0.01 |

**Table S2. Model synaptic weights for abstract rule processing. Related to Figures 7 and S13.**

|  | **Synaptic Weights** |
| --- | --- |
| PFC middle to PFC concrete rule | 0.02 |
| PFC concrete rule within chains | 0.0012 |
| PFC concrete rule to MD | 0.0009 |
| PFC choice to MD | 0.0009 |
| PFC choice within chains | 0.002 |
| MD to PFC choice | 0.0036 |
| MD to PFC concrete rule | 0.012 |
| MD to PFC inhibitory cells | 0.001 |
| PFC abstract rule to inhibitory cells | 0.005 |
| Inhibitory cells to PFC concrete rule | 0.5 |
| Inhibitory cells to PFC choice | 0.0025 |

**Table S3. Model synaptic weights for concrete rule and choice processing. Related to Figures 7 and S13.**
